# A genetically encoded biosensor reveals heterogenous cAMP dynamics coordinating growth and resuscitation in Mycobacterium tuberculosis

**DOI:** 10.64898/2026.06.02.729633

**Authors:** Shaun Wachter, Srivathsa Shankar Kurpad, Amit Singh, Neeraj Dhar

## Abstract

3′,5′- Cyclic adenosine monophosphate (cAMP) serves as a global regulatory hub in *Mycobacterium tuberculosis* (Mtb), which dedicates approximately 1% of its genome to the synthesis, degradation and sensing of this secondary messenger. Despite cAMP’s central role in Mtb virulence and metabolic adaptation, we currently do not have any tools to monitor cAMP dynamics in real time at single-cell resolution. Here, we report the development and validation of rmgCarvi, a genetically encoded biosensor adapted from eukaryotic systems for use in mycobacteria. By utilizing cAMP-responsive cpGFP and constitutive mCherry fluorescence, rmgCarvi provides a high-fidelity ratiometric readout of intracellular cAMP levels, revealing that carbon source and growth state act as primary determinants of cAMP dynamics. Real-time single-cell imaging using rmgCarvi demonstrated robust correlations between cAMP and growth rate, with pronounced cAMP spikes coinciding with cell division. In starved Mtb populations, a bimodal distribution in cAMP levels dictates a distinct hysteresis during regrowth upon nutrient supplementation. Low-cAMP subpopulations are associated with strong induction in cAMP levels and a longer lag before initiation of growth, while high cAMP cells do not exhibit significant induction in cAMP and have recovery times uncorrelated with prior cAMP levels. Finally, use of rmgCarvi in infection studies captured host species-specific responses, including a strong cAMP induction upon internalization followed by a decline in murine, but not human, macrophages. By resolving cAMP dynamics with unprecedented cellular and temporal resolution, rmgCarvi provides a framework for understanding how Mtb tunes this crucial metabolite to navigate the complex microenvironments in the host.

**Significance Statement:** We report the evaluation of a ratiometric genetic biosensor, rmgCarvi, for real-time single-cell monitoring of cAMP dynamics in mycobacteria. This biosensor reveals how *Mycobacterium tuberculosis* precisely tunes cAMP within a narrow range (“Goldilocks zone”) to coordinate metabolic adaptation, and growth across complex nutrient environments. In nutrient-starved populations, we identify a bimodal cAMP distribution that functions as a phenotypic memory, dictating resuscitation kinetics upon nutrient restoration. As the first cAMP biosensor validated in any mycobacterial species, rmgCarvi facilitates the discovery of host-specific signaling dynamics during infection and provides a platform for screening of compounds that perturb cAMP homeostasis. This study provides a framework to explore how Mtb utilizes this secondary messenger molecule to survive and adapt within complex host environments.

## Introduction

The success of *Mycobacterium tuberculosis* (Mtb) as an obligate human pathogen is due to its extraordinary ability to sense and adapt to the diverse, stressful microenvironments of the host (1, 2). Central to this adaptation is a complex signaling network governed by nucleotide secondary messengers, which relay environmental cues to downstream effectors to reprogram physiology (3, 4). Among these, 3′,5′- cyclic adenosine monophosphate (cAMP) serves as a primary global regulatory hub, regulating a variety of Mtb processes necessary for carbon source utilization, adaptation to the host environment, and establishment of the Mtb intra-phagosomal niche, amongst others (3–10). Mtb dedicates approximately 1% of its coding capacity to cAMP signaling, encoding an unusually large repertoire of at least 16 adenylyl cyclases (ACs) to synthesize the molecule and a wide array of cAMP-binding effector proteins (4, 7, 8). Among the 16 ACs encoded by the Mtb genome, Rv3645 (MacE) appears to be the main source of cAMP during standard axenic growth (8). Conversely, AC Rv0386 plays a key role in establishing the intra-phagosomal niche (9), and AC Rv2212 produces cAMP under acidic, fatty-acid–rich conditions and has been implicated in resuscitation of Mtb from dormancy (10, 11). Mtb also encodes two phosphodiesterase enzymes (PDE), Rv1339 and Rv0805, which degrades cAMP to adenylyl monophosphate to regulate cAMP content (12, 13).

Unlike the classical model in *Escherichia coli*, where cAMP primarily mediates catabolite repression through a single adenylyl cyclase (14, 15), Mtb utilizes its diverse adenylyl cyclases to integrate a multitude of host-associated signals, including pH, fatty acids, and CO_2_ levels (3–10, 16). Intracellularly, cAMP modulates Mtb gene expression notably through the cAMP-receptor protein (CRP) family and protein function (16–19), including the post-translational regulation of metabolic enzymes via lysine acetylation (20). Beyond its internal roles, Mtb is distinguished by its ability to secrete massive amounts of cAMP into host macrophages (9, 21). This “cAMP intoxication” subverts host innate immunity by elevating intramacrophage cAMP levels, thereby inhibiting inflammatory cytokine secretion and potentially arresting phagosome maturation (9, 16, 22). Crucially, it is not merely the basal abundance of cAMP but rather Mtb’s ability to dynamically modulate cAMP levels in response to environmental cues that appears to underlie its adaptive success (3, 6, 8–11, 16, 20, 22). Despite cAMP’s established role in Mtb virulence and metabolic adaptation, reflected in the vast repertoire of genes involved in its synthesis and interactions, our understanding of its dynamic regulation remains limited. Current approaches rely on genetic dissection of individual enzymes and population-level invasive biochemical assays (mass spectrometry, ELISA) that provide only steady-state snapshots in bulk samples, obscuring single-cell heterogeneity or temporal dynamics. Genetically encoded biosensors offer a solution by enabling non-invasive, real-time monitoring of metabolite fluctuations at single-cell resolution and could thus provide a better mechanistic understanding of cAMP signaling in Mtb biology (23).

In this study, we address this challenge by developing and characterizing rmgCarvi, a genetically encoded, ratiometric cAMP biosensor optimized for mycobacteria. Building upon gCarvi, a eukaryotic-based tool (24), this biosensor utilizes cAMP-responsive cpGFP and constitutive mCherry fluorescence that enables high cellular and temporal measurements of cAMP dynamics at the single-cell level using flow cytometry or live-cell microscopy. The biosensor was validated in *Mycobacterium smegmatis* (Msm) and Mtb and revealed carbon source-dependent and species-specific differences in cAMP responses upon exposure to stresses. Single-cell imaging of Mtb expressing rmgCarvi revealed that growth rates closely correlated with cAMP content, with marked increase in cAMP levels detected during cell division. Nutrient-starved Mtb populations exhibited bimodal phenotypic bifurcation into low and high-cAMP populations that displayed contrasting regrowth kinetics upon nutrient restoration, suggestive of hysteresis in cAMP levels.

Application of the rmgCarvi biosensor during macrophage infection uncovered host species-specific differences in cAMP dynamics. Beyond these mechanistic insights, rmgCarvi also enabled high-throughput screening of compounds that modulate cAMP levels, offering a scalable platform for identifying novel anti-TB therapeutics. Our findings reveal that Mtb maintains cAMP levels within a narrow growth-permissive range, a “Goldilocks zone”, that is dependent on the carbon source or environmental condition. We envision rmgCarvi becoming an essential tool for the broader microbiology community, facilitating systematic dissection of cAMP signaling across diverse bacterial pathogens and revealing conserved principles of metabolite-dependent regulation.

## Results

### rmgCarvi is a functional ratiometric cAMP reporter in mycobacteria

Several genetically encoded biosensors have been developed for capturing cAMP dynamics in eukaryotic cells (24–29). However, to our knowledge none have been developed to monitor cAMP levels in mycobacteria. To address this shortcoming, we cloned and expressed several existing cAMP biosensors in Msm, each of which failed either due to weak fluorescence (e.g. Pink Flamindo (25)) or suspected toxicity (e.g. cAMPFIRE (26)). Most of these biosensors utilize the cAMP-binding domains (CBDs) of eukaryotic guanine nucleotide exchange proteins (Epac-1 and Epac-2) or protein kinase A (25–29).

A recent study reported the construction of a highly specific ratiometric monomeric cAMP indicator, gCarvi, that utilizes the CBD of the *E. coli* cAMP receptor protein (CRP) (24). gCarvi possesses the key features required for a functional mycobacterial cAMP reporter - is small and monomeric, uses a single fluorophore (cpGFP) for cAMP sensing, offers a ratiometric readout (cpGFP:mCherry) to account for cell-to-cell heterogeneity, exhibits high specificity for cAMP, a high dynamic range and sensitivity to account for fluctuations in cAMP levels and crucially incorporates a CBD of bacterial origin. The CBD in gCarvi comprises of *E. coli’s* CRP N-terminal cyclic nucleotide binding module and its long a-helix (C-helix) fused to cpGFP at the C-terminus and mCherry at the N-terminus (24). Binding of cAMP results in changes in fluorescence intensity on the green channel with the red channel serving as an internal control. Ratiometric gCarvi was cloned for expression in mycobacteria as a fusion with six N-terminal codons from the highly expressed Mtb *hsp65* gene (*rv0440*) to enhance ribosome loading in a mycobacterial background (rmgCarvi) (30) (**Fig. 1A**). rmgCarvi was cloned and expressed using a single-copy chromosomal integrative plasmid (rmgCarvi_int_) or using a multicopy episomal plasmid (rmgCarvi_epi_) in Mtb and Msm (**Table S1)**. Expression of rmgCarvi did not affect the growth of Mtb whether cultured in standard mycobacterial growth media (7H9+OADC) or in media containing fatty acids and cholesterol as the carbon source (BCP) (**Fig. S1A and B**). rmgCarvi production was validated by Western blot analysis (**Fig. 1B, Fig. S1C**). As expected, Mtb-rmgCarvi_epi_ expressed ∼4.7 fold more protein than Mtb-rmgCarvi_int_, during exponential phase, with levels further increasing in stationary phase. As reported previously for gCarvi (24), expression of rmgCarvi in Mtb showed strong excitation peaks at 500 nm and 585 nm and emission peaks at 525 nm and 622 nm for cpGFP and mCherry respectively, when compared to wild-type Mtb (**Fig. 1C, D**). To evaluate the dose response and functionality of the reporter, bacterial lysates from strains expressing rmgCarvi_int_ and rmgCarvi_epi_ were exposed to different concentrations of cAMP (**Fig. 1E, F**). rmgCarvi was able to detect cAMP in the range (0.5-100 μM) with a dynamic range of ∼4 in case of rmgCarvi_int_ and ∼12 in case of rmgCarvi_epi_ (**Fig. 1E**). As expected, mCherry fluorescence was non-responsive to cAMP concentrations and can therefore function as an internal normalizing control in the ratiometric probe (**Fig. 1F**).

**Figure 1.**
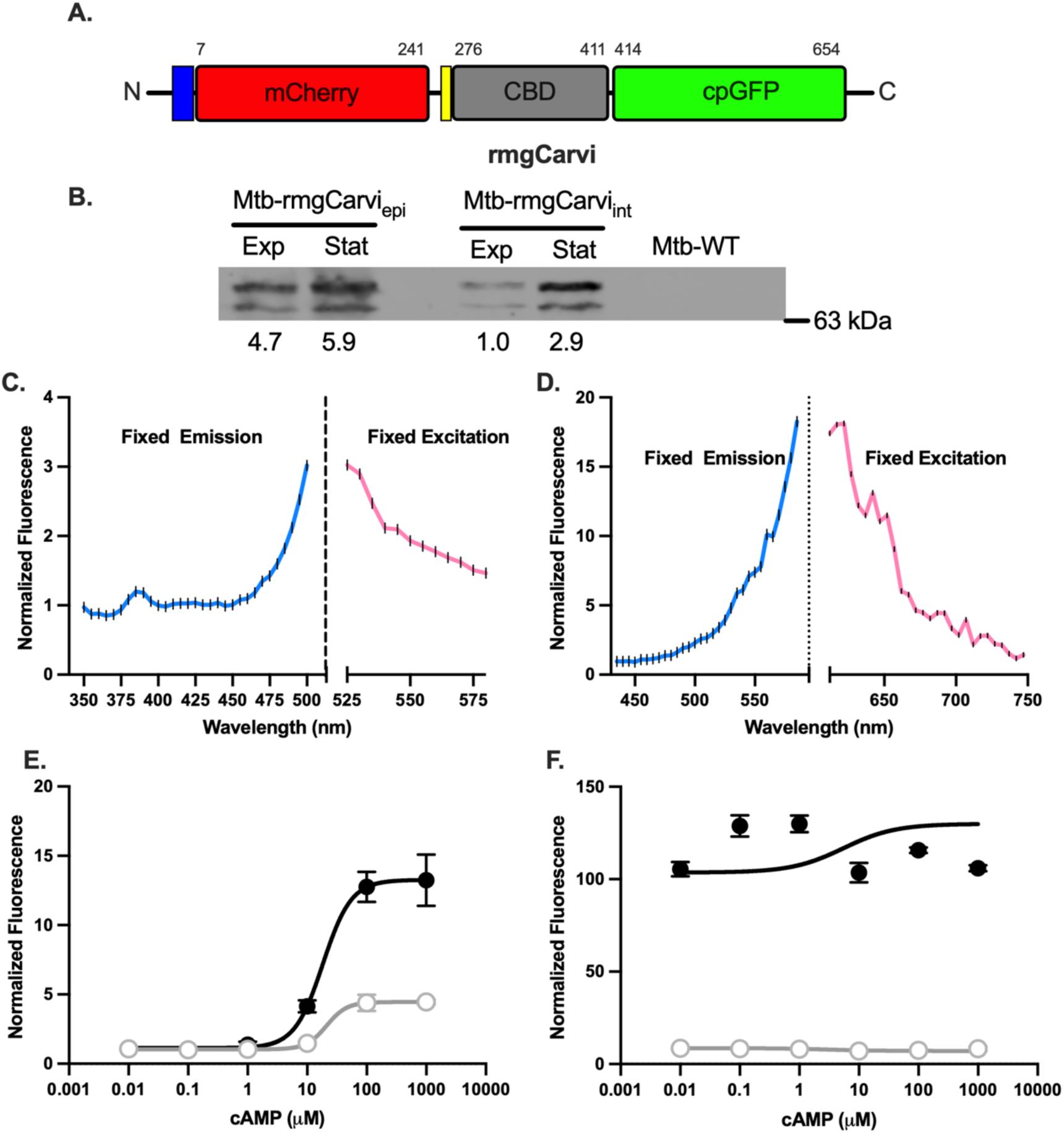
Characterization of rmgCarvi as a ratiometric cAMP biosensor in mycobacteria. (**A**) Schematic of the rmgCarvi construct. The *E. coli* CBD is flanked by mCherry at the N-terminus and cpGFP at the C-terminus and was fused to the first 6 amino acids of Mtb *hsp65* gene (blue box). The location of the FLAG-tag is indicated by a yellow box. The numbers on top represent the amino acid numbers. (**B**) Immunoblots depicting the expression of rmgCarvi in Mtb mc^2^6206, from either an episomal (rmgCarvi_epi_) or an integrative (rmgCarvi_int_) plasmid in cells harvested from cultures in exponential (Exp) or stationary (Stat) phase, probed with an anti-FLAG antibody. The numbers below indicate the fold expression normalized to total protein and to levels in exponential cultures of Mtb-rmgCarvi_int_. (**C, D**) Excitation (blue) and emission (magenta) spectra of Mtb-rmgCarvi_epi_ for detection of cpGFP (500 nm excitation and 525 nm emission) (**C)**, and mCherry (587 nm excitation and 612 nm emission) (**D**). Spectra were normalized to the values obtained from parental wild-type Mtb. (**E, F**) Dose-response curves obtained by exposing bacterial lysates from Msm-rmgCarvi_epi_ (filled circles) and Msm-rmgCarvi_int_ (empty circles) to different concentrations of cAMP and measuring cpGFP (**E**) or mCherry (**F**) fluorescence. Fold-change and dynamic ranges were obtained by first subtracting fluorescence values from the wild type Msm strain and then normalizing to *F_min_*. mCherry fluorescence is insensitive to changes in cAMP content. N=3, Symbols represent the mean ± SD.

### rmgCarvi accurately tracks cAMP levels in mycobacteria

*M. smegmatis* and *M. tuberculosis* strains expressing rmgCarvi_epi_ were cultured in 7H9 base medium supplemented with either ADS or OADC or without any carbon source. Analysis of the ratiometric rmgCarvi fluorescence values suggests that Msm has higher cAMP levels than Mtb, as reported previously (31). Interestingly, the presence of fatty acids in the growth media (7H9+OADC) did not significantly alter cAMP levels in Msm or Mtb during exponential growth (**Fig. 2A, B**). Additionally, regardless of the carbon source, cAMP levels in both Msm and Mtb were significantly reduced in stationary phase (**Fig. S2A, B**). This was despite the fact that Mtb in stationary-phase cultures contained significantly higher amount of rmgCarvi than bacteria in exponential phase (**Fig. 1B**), further supporting biosensor functionality. We observed that cAMP levels in Msm were less sensitive to carbon source than Mtb (**Fig. S2A, B**). cAMP is a central mediator of fatty acid metabolism in Mtb (8, 20), and earlier studies have demonstrated an inverse relationship between cAMP levels and ability of Mtb to utilize cholesterol (32–35). As expected, Mtb grown in presence of cholesterol (BCP) exhibited very low rmgCarvi fluorescence ratios, an effect not observed in case of Msm (**Fig. 2A, B)**. Introduction of an R324A mutation in the cAMP binding domain of rmgCarvi, that was previously shown to disrupt cAMP binding (36), rendered rmgCarvi non-responsive to cAMP levels in mycobacteria (**Fig. 2A**). Exposure of Mtb-rmgCarvi_epi_ to GSK2556286, a known adenylyl cyclase agonist, significantly increased rmgCarvi fluorescence values in Mtb, but only in bacteria cultured in cholesterol containing medium (BCP), as has been reported previously (35) (**Fig. 2C**, *p* < 0.001). These observations were further confirmed biochemically by measuring cAMP levels using a commercial cAMP enzyme-linked immunosorbent assay (ELISA) kit (**Fig. 2D**, *p* < 0.005). Finally, to validate the functionality of rmgCarvi reporter, we adopted a genetic approach to perturb mycobacterial cAMP levels. cAMP is synthesized from ATP by adenylyl cyclases and is hydrolyzed and degraded by phosphodiesterase enzymes. The Mtb genome encodes for ∼16 distinct adenylyl cyclase genes (4), among which *rv3645* (*macE*) was recently shown to be the dominant source of cAMP, in standard lab media (8). Mtb also encodes for two phosphodiesterases, *rv0805* and *rv1339* (12, 31). We used CRISPRi (37) to construct genetic knockdown strains of *rv3645*, *rv0805*, and *rv1339*. The CRISPRi strains were grown to early exponential phase in 7H9+ADS and the target genes were knocked down by exposure to 100 ng/mL anhydrotetracycline (aTc) for 48 h. qRT-PCR analyses confirmed ∼75-85% reduction in expression of the targeted genes in comparison to the no aTc control (**Fig. S3**). Knockdown of *rv3645* led to significant decrease in rmgCarvi fluorescence levels, consistent with its role as the major adenylyl cyclase in Mtb. In case of the two phosphodiesterases, while knockdown of *rv1339* resulted in a considerable increase in rmgCarvi fluorescence, knockdown of *rv0805* did not cause any significant difference in fluorescence ratios (**Fig. 2E**). Again, these results were confirmed by measurement of cAMP levels using the conventional cAMP ELISA kit (**Fig. 2F**). We conclude that rmgCarvi functions as a functional cAMP reporter in both Msm and Mtb, and that the cpGFP:mCherry ratiometric fluorescence measurements are a consistent and reliable proxy for cAMP content.

**Figure 2.**
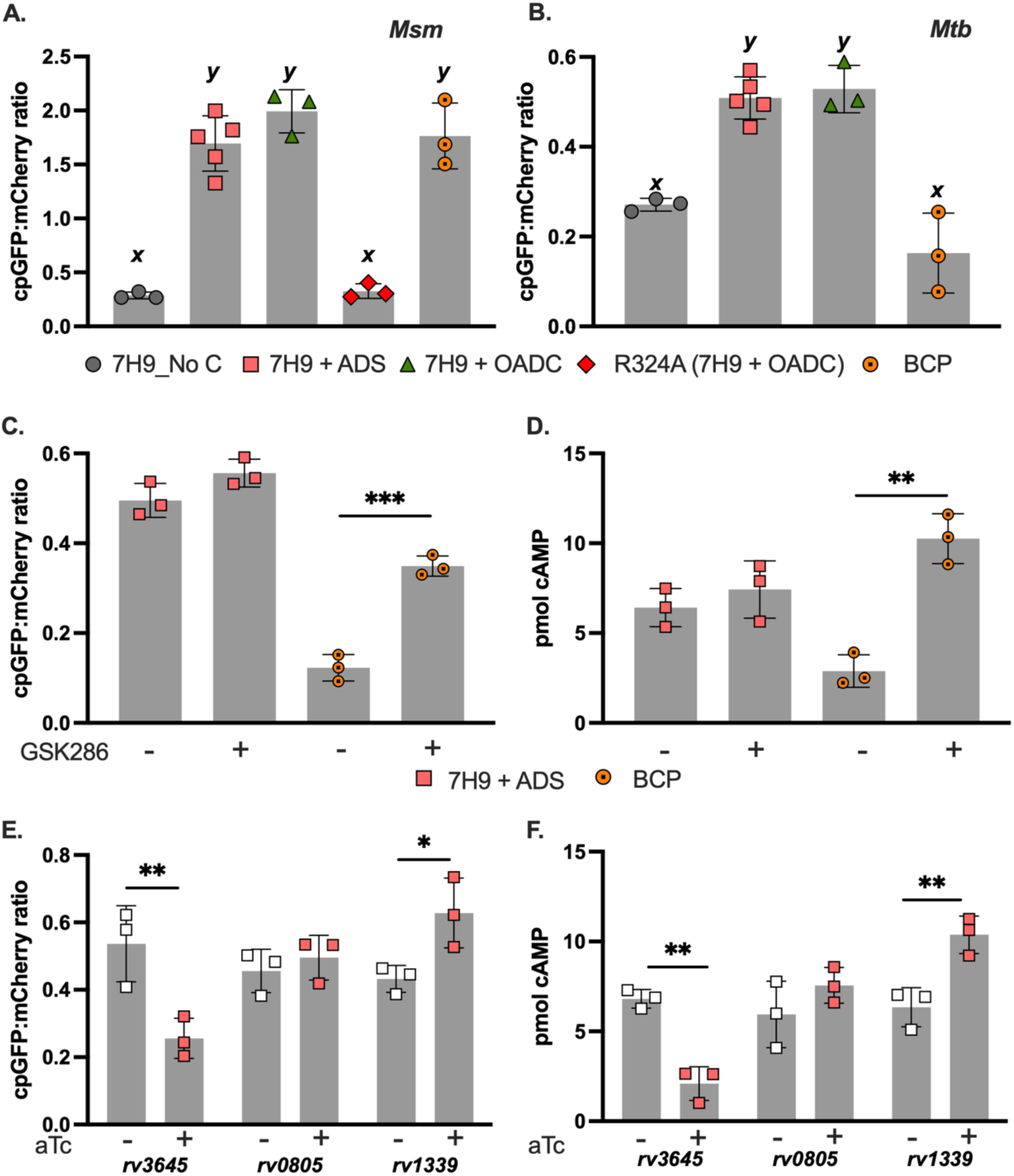
rmgCarvi accurately reports cAMP levels in mycobacteria. (**A, B**) Bacteria from exponential cultures of Msm-rmgCarvi_epi_ or Mtb-rmgCarvi_epi_ was sub-cultured into various growth media - 7H9-no carbon source (7H9_No C, gray circles); 7H9+ADS (orange squares); 7H9+OADC (green triangles) or cholesterol based BCP media (orange dotted circle). Ratiometric rmgCarvi values (*y-axis*) shows that presence of cholesterol causes significant decrease in cAMP levels in Mtb but not in Msm. Introduction of the R324A mutation into rmgCarvi’s CBD domain abolishes rmgCarvi’s ability to respond to cAMP levels (red diamond). N=3-5; and the characters above the bars represent the different groupings determined by ordinary one-way ANOVA. (**C, D**) Exposure of Mtb-rmgCarvi_epi_ (mc^2^6206 strain) cultures to a known adenylyl cyclase agonist, GSK286, results in increased cAMP levels only when grown in media containing cholesterol, measured by ratiometric rmgCarvi fluorescence (**C**) or using a commercial cAMP ELISA kit (**D**). N=3. (**E, F**) CRISPRi-mediated knock-down of cAMP-related genes significantly alters cAMP content in Mtb-rmgCarvi_epi_ (mc^2^6206 strain) measured by ratiometric rmgCarvi fluorescence (**E**), confirmed by cAMP ELISA assay (**F**). N=3. All data represent the means ± SD. Unpaired t-test, * - *p* < 0.05, ** - *p* < 0.01, *** - *p* < 0.001.

### Growth phase and carbon source govern mycobacterial cAMP levels

Upon validation of rmgCarvi, we explored its potential to provide a readout of intrabacterial cAMP levels in a dynamic manner. Msm and Mtb transformants expressing rmgCarvi_epi_ were cultured in standard growth media with or without supplementation of fatty acid (oleic acid) or cholesterol (BCP). They were also exposed to acidic pH or high salt conditions, both of which have been previously shown to alter cAMP levels in Mtb (16, 38). Exposure to acidic pH (pH 5.5) or physiological intramacrophage NaCl concentrations (∼250 mM), induced cAMP levels in Mtb but not in Msm (**Fig. 3A, Fig. S4)**.

**Figure 3.**
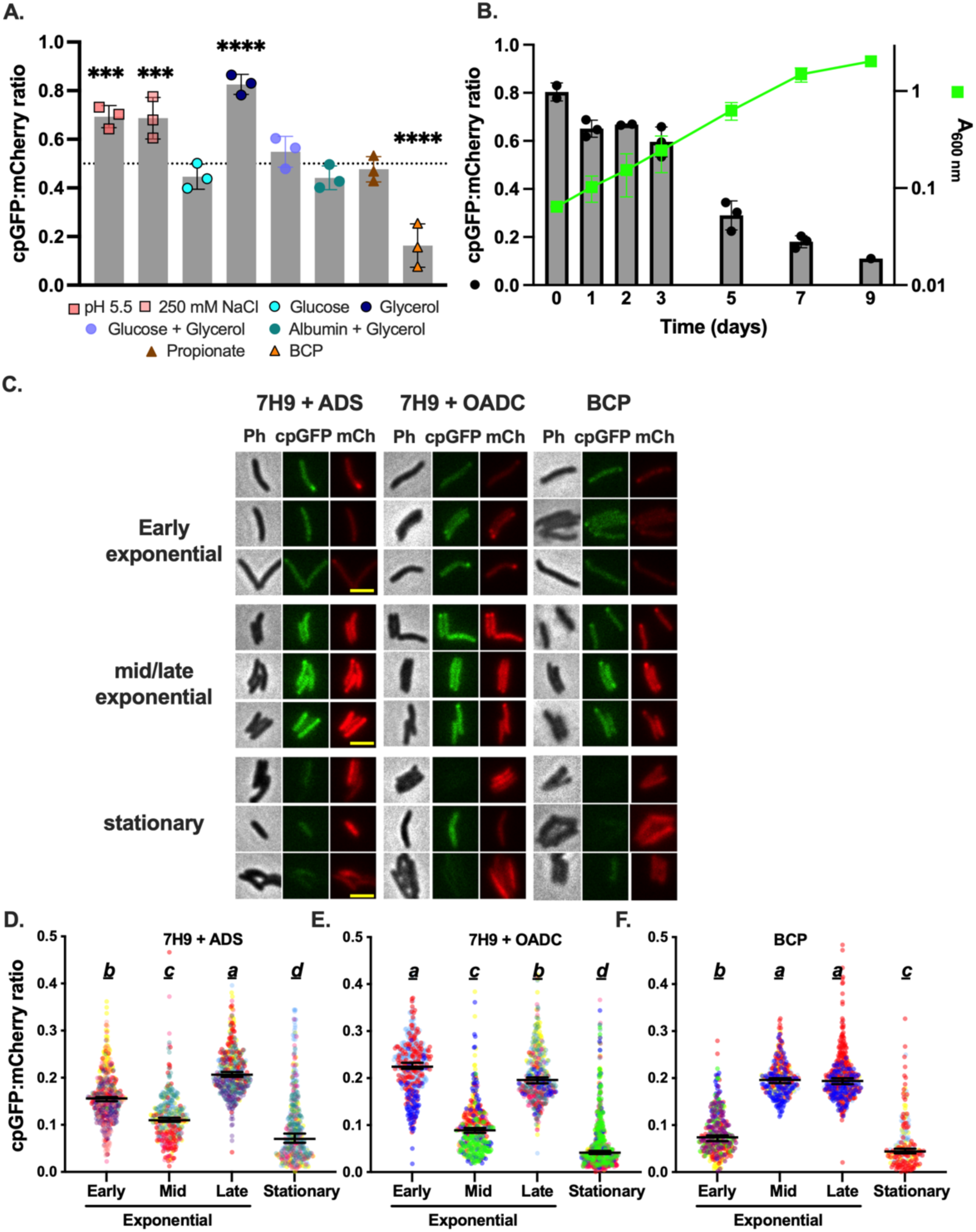
Ratiometric rmgCarvi reveals growth state and carbon source dependent changes in cAMP levels in *M. tuberculosis*. (**A**) Mtb-rmgCarvi_epi_ (mc^2^6206 strain) was cultured in 7H9-based growth media, supplemented with various carbon sources or exposed to acidic pH (pH 5.5) or high salt concentrations (250 mM NaCl) and ratiometric rmgCarvi fluorescence measured. The dotted horizontal line indicates the fluorescence values in bacteria cultured in 7H9+ADS (containing glucose and glycerol) (Fig. 2B). N=3. Data were analyzed by Ordinary one-way ANOVA with Dunnett’s multiple comparison test. *** - *p* < 0.001; **** - *p* < 0.0001 (**B**) Plot depicts the ratiometric rmgCarvi fluorescence (left *y-axis*) and A_600 nm_ (right *y-axis*) of Mtb-rmgCarvi_epi_ (mc^2^6206 strain) strain cultured in 7H9+ADS. Measurements were made at the indicated time points during the different phases of growth. N=3. (**C-F**) Representative snapshots of Mtb-rmgCarvi_epi_ (Erdman strain) bacteria cultured in 7H9+ADS or 7H9+OADC or BCP from different phases of growth. The phase (Ph), cpGFP, and mCherry (mCh) channels are shown. Scale bar represents 3 μm. More images are provided in **Fig. S6**. (**D**) Cultures were classified as early exponential (A_600 nm_ 0.1-0.3 for 7H9+ADS and 7H9+OADC; 0.1-0.2 for BCP), mid-exponential (A_600 nm_ 0.3-0.5 for 7H9 ADS and 7H9 OADC; 0.2-0.4 for BCP), late-exponential (A_600 nm_ 0.5-1 for 7H9 ADS and 7H9 OADC; 0.4-0.7 for BCP), and stationary (>1 for 7H9 ADS and 7H9 OADC; >0.7 for BCP). Acquired images were analyzed using FIJI and fluorescence intensity values plotted in (**D-F**). Each dot represents a single cell. Biological replicates are color-coded, and descriptive statistics of the different populations have been provided in **Table S2**. Every growth phase dataset in each media represents two to four biological replicates and 200-600 individual cells analyzed under each condition. The underlined characters above the scatter plots represent the different groupings determined by Brown-Forsythe and Welch ANOVA with Games-Howell’s multiple comparisons test. Data in (**A, B**) represent the means ± SD. Data in (**D-F**) represent the medians ± 95% CI.

As standard mycobacterial growth media contains a mixture of carbon sources, we decided to evaluate the contribution of each individual carbon source to cAMP levels in Mtb. We found glycerol on its own caused a significant increase in cAMP levels, while supplementation with glucose or albumin prevented this increase (**Fig. 3A**). This is suggestive of a mechanism like the *E. coli* cAMP-CRP carbon choice paradigm, although unlike *E. coli*, Mtb has been shown to co-metabolize multiple carbon sources (14, 39, 40). Like oleic acid (**Fig. 2B)**, the use of propionate as a carbon source did not cause any significant difference in cAMP levels, also confirming that the drastic decrease in cAMP levels seen in case of BCP media was likely driven by cholesterol. These results highlight the strong influence of carbon source on cAMP levels in Mtb, but not in Msm.

We cultured the mycobacterial strains in standard 7H9 media containing OADC and analyzed bacteria at different time points during their growth using a plate reader or by flow cytometry (**Fig. 3B, Fig. S5**). In both Msm and Mtb, subculture into fresh medium was accompanied by high cAMP levels which reduced and stabilized during mid-exponential phase and then further dropped as the cultures approached stationary phase. To better capture rmgCarvi fluorescence and cAMP heterogeneity at the single-cell level during axenic mycobacterial growth, we cultured Mtb-rmgCarvi_epi_ in the different media formulations (7H9+ADS, 7H9+OADC and BCP) and imaged the cells periodically during the different phases of growth using a fluorescent microscope (**Fig. 3C, Fig. S6**). We carried out quantitative analysis of ratiometric rmgCarvi fluorescence in Mtb across exponential (early, mid, late) and stationary phase. Interestingly, in bacteria grown in 7H9+ADS and 7H9+OADC media, cAMP levels decreased during the transition from early to mid-exponential phase but increased during the transition from mid-to late-exponential phase (**Fig. 3D, E**). An alternate trend was observed in bacteria cultured in BCP media, where fatty acids and cholesterol are the main carbon sources. The cAMP levels increased as cultures aged from early to mid-exponential, stayed high even in late-exponential phase, and only dropped during stationary phase (**Fig. 3F**). Considering the relationship between Mtb cAMP and cholesterol/fatty acid metabolism, this may reflect a slow depletion of these carbon sources in culture, facilitating a gradual increase in cAMP levels (8, 10, 34). Regardless of the culture condition, single-cell heterogeneity in cAMP levels (using CV^2^ as a metric) increased in Mtb cultures as they transitioned to stationary phase (**Fig. 3D-F, Table S2**). This relationship between cAMP and growth phase of the bacterial culture underlies the dynamic metabolic flexibility imparted by cAMP and highlights the need to specify culture density when reporting cAMP levels in mycobacteria. More importantly, this tool allows us to capture cAMP dynamics with single-cell resolution.

### Single-cell imaging of rmgCarvi in *M. tuberculosis* unveils cAMP dynamics during growth and division

The results so far suggest that the intracellular levels of cAMP in Mtb are dynamically modulated in response to the growth phase and the available carbon source. This further highlights the value of a biosensor like rmgCarvi, which captures cAMP production dynamics rather than snapshots provided by conventional endpoint biochemical assays. To explore cAMP dynamics in Mtb at the single-cell level in real-time, we utilized a microfluidics-based microscopy platform in combination with automated time-lapse microscopy (41). Mtb expressing rmgCarvi_int_ was cultured in 7H9+OADC media to early exponential phase, and cells seeded into a microfluidic device for time-lapse imaging. Using this setup, we were able to capture dynamic fluctuations in cAMP in individual bacteria which appeared to correspond to periods of growth during its cell division cycle (**Fig. 4A**). To detect if indeed cAMP levels correlate with growth capacity, we analyzed a subset of cells with varying growth rates (N = 165, across four independent experiments, median growth rate 0.024 h^-1^, range 0.006-0.06 h^-1^). We noted moderate positive correlations between cAMP content and growth rate (**Fig. 4B**, Pearson correlation coefficient *r* = 0.44, *p* < 0.0001) and cAMP content and growth velocity (**Fig. 4C**, Pearson correlation coefficient *r* = 0.37, *p* < 0.0001) in Mtb cells grown under these experimental conditions. These moderate correlations were largely reflective of the differences in culture history across the four independent experiments, as this correlation was much stronger when analyzing the cells within each experimental replicate separately (**Fig. S7**). These observations are consistent with our earlier findings where we found that slight differences in culture condition or culture history can significantly influence cAMP content. Interestingly, when analyzing these single cells for correlations to growth rate, we noted a phenomenon where cAMP levels appeared to spike briefly around the time when the cell was getting ready to divide (**Fig. 4A**). As we didn’t have a fluorescently tagged marker protein to precisely identify the time of cell division, we chose to use the appearance of furrow near the septal region, on the phase channel. Upon aligning Mtb cells based on the appearance of this marker, we observed a strong spike in cAMP levels approximately 2 hours prior to septum furrow appearance (**Fig. 4D, E**). Importantly, this pulsing was detected only on the cpGFP channel, which reports on cAMP content, and not on the mCherry channel (**Fig. 4E, Fig. S8**). Similar results were obtained when culturing Mtb in 7H9+ADS media or upon imaging the Mtb-rmgCarvi_epi_ strain. These observations indicate that cAMP signaling likely plays a key role in Mtb cytokinesis, although further work is required to elucidate the precise mechanism.

**Figure 4.**
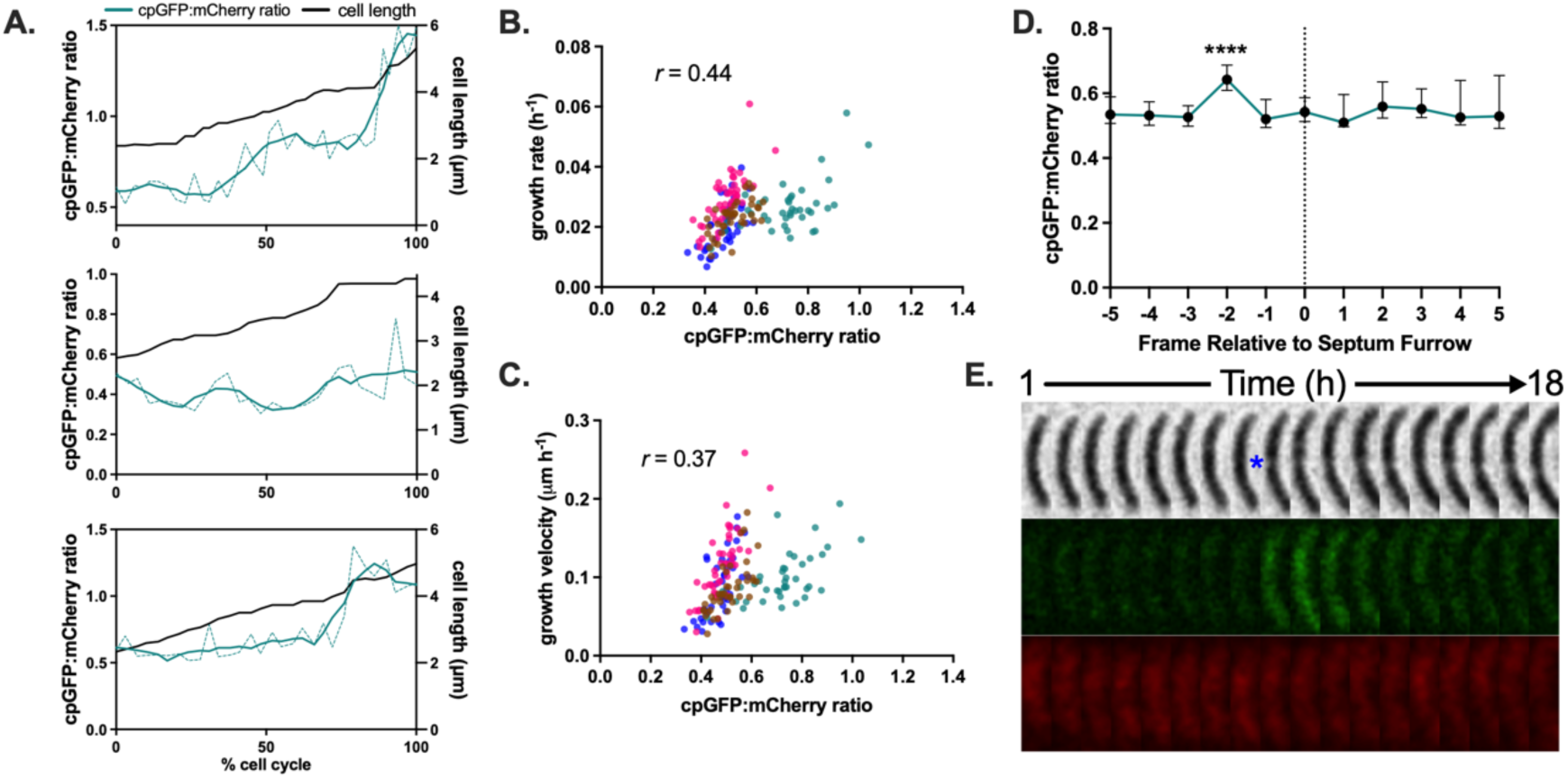
Time-lapse microscopy reveals cAMP dynamics during *M. tuberculosis* growth and division cycles. (**A**) Plots depict the ratiometric rmgCarvi fluorescence (solid green lines – smoothed data, dashed green lines – raw data, left *y-axis*) and cell-length (black lines, right *y-axis*) of representative Mtb-rmgCarvi_int_ (Erdman) bacteria, mapped from their birth to division (*x-axis*). (**B, C**) Images of Mtb-rmgCarvi_int_ bacteria, growing and dividing inside the microfluidic device, were acquired every hour, across four independent experiments. For each cell observed, ratiometric rmgCarvi fluorescence during the growth period was measured and plotted against its growth rate (**B**) or growth velocity (**C**). Plot depicts the correlation with the Pearson correlation coefficient (*r*) indicated, *p* < 0.0001. Each dot represents an individual cell, and the four biological replicates are color-coded (N=165). Plots for each replicate set are provided in **Fig. S7**. (**D**) Ratiometric rmgCarvi fluorescence in Mtb-rmgCarvi_int_ bacteria, plotted five frames prior and post formation of septum furrow. N=81, across three independent experiments. Data represent medians ± 95% CI. *** - *p* < 0.0001, One-way ANOVA with Geisser-Greenhouse and Holm-Sidak’s corrections. (**E**) Representative image sequence of an Mtb-rmgCarvi_int_ bacterial cell before, and after cell division, depicting the cAMP pulsing event. The appearance of the furrow in the septal region is indicated with a blue asterisk. Bacteria were cultured in a microfluidic device and images acquired on phase (top), cpGFP (Ex_475 nm_/Em_535 nm_, middle) and mCherry (Ex_575 nm_/Em_642 nm_, bottom) channels using a 100X objective. The numbers above indicate the time span (h) of the image series.

### Phenotypic bifurcation and hysteresis drive reawakening from dormancy in *M*. tuberculosis

Several studies have implicated cAMP to play an important role in resuscitation of Mtb from dormancy or from a metabolic inactive state (11, 42, 43). Recovery of dormant Mtb was also shown to be stimulated by unsaturated fatty acids through the action of a fatty acid-activated adenylyl cyclase (10, 11, 42, 44). To mimic growth arrest and a metabolic inactive state in Mtb, we employed an extended starvation model (45), in which Mtb-rmgCarvi_epi_ cells were starved for six weeks in PBS+0.02% tyloxapol. Flow cytometry analysis of cAMP levels during recovery in either 7H9+ADS or 7H9+OADC, revealed a significant induction of cAMP, independent of long-chain unsaturated fatty acids **(Fig. 5A)**. To better capture cAMP dynamics and response during recovery from starvation, we carried out time-lapse imaging on starved Mtb-rmgCarvi_int_ cultures. Bacteria from cultures starved for five to six weeks in PBS + tyloxapol, were seeded into the microfluidics device and subsequently exposed to standard 7H9+OADC growth media. We noted a clear bi-modal distribution of rmgCarvi fluorescence amongst the starved cells representing low and high cAMP containing Mtb subpopulations (N=150 across two independent experiments) (**Fig. 5B, Fig. S9**). This was rather surprising as it suggests that high levels of cAMP are not invariably associated with active growth and may be insufficient to sustain it. Upon exposure to fresh growth medium (7H9+OADC), we detected disparate responses in the two subpopulations. While the low cAMP population exhibited a strong induction in cAMP levels, the high cAMP population responded more heterogeneously and did not show significant increase in cAMP levels (**Fig. S10A, B**). Indeed, we found a strong inverse correlation between cAMP content during starvation and the magnitude of cAMP induction during recovery from starvation, perhaps indicating some cAMP threshold that must be met before growth can resume (**Fig. 5C**, R^2^ = 0.97). However, interestingly, we also invariably found that regrowth only occurred after reduction of cAMP content after the initial cAMP burst (**Fig. 5A**). Consequently, in the low cAMP population, although cAMP induction occurs, growth did not resume in majority of the cells during the observation period by time-lapse microscopy (∼169 hours). Of the ∼2000 Mtb cells imaged, only 90 recovered growth (<5% recovery). Amongst the regrowing bacteria, the lag time prior to regrowth was strongly inversely correlated to the cAMP content during starvation, however only in case of cells in the low cAMP population (**Fig. 5D**, Pearson correlation coefficient, *r* =-0.77, *p* < 0.0001). Amongst the high cAMP population, there was no correlation between cAMP levels and the lag-time before recovery from starvation (**Fig. S10C**, Pearson correlation coefficient, *r* =-0.1, *p* = 0.45; low and high combined data plotted in **Fig. S10D**, Pearson correlation coefficient, *r* = - 0.35, *p* = 0.0004). Additionally, in cells where regrowth did occur during the timeframe of the experiment, growth rate and growth velocity was again positively correlated to cAMP content, however the strength of this correlation differs depending on whether the cell belonged to the low or high cAMP sub-populations during starvation (**Fig. 5E, F and Fig. S10E, F**). In summary, low [cAMP] during starvation leads to a strong induction of cAMP during recovery, resulting in a longer but predictable lag between exposure to growth-permissive conditions and regrowth. High [cAMP] during starvation leads to a smaller magnitude of cAMP induction, but the timing of re-growth is not linked to cAMP content. This would suggest that heterogeneity in cAMP levels functions as a phenotypic memory that dictates the timing of re-growth following extended starvation and that certain Mtb cells continue to maintain high cAMP levels that allow them to recover more rapidly during resuscitation (**Fig. S10D**).

**Figure 5.**
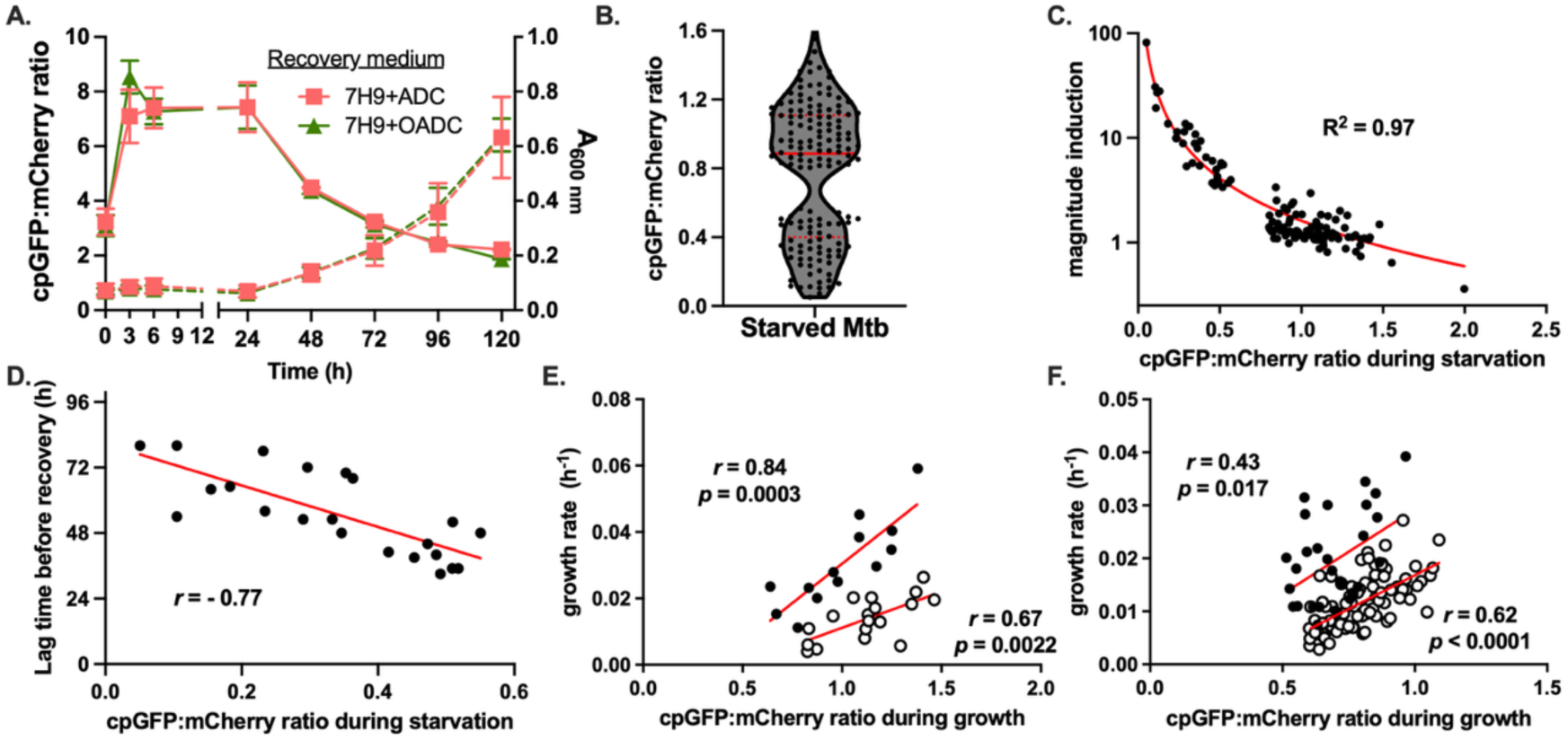
Bimodality and hysteresis in cAMP levels during dormancy and regrowth in *M. tuberculosis*. (**A**) Mtb-rmgCarvi_epi_ (Erdman) starved for 6-8 weeks in PBS + 0.02% tyloxapol, were recovered in 7H9+ADS (red line) or 7H9+OADC (green line) (*left y-axis*) and fluorescence ratios measured by flow cytometry and growth monitored (A_600 nm_, dotted lines *right y-axis*) at indicated time points, over 120 h (*x-axis*) (N=3, 10000 events per sample). Data represents means ± SD (**B**) Measurement of ratiometric rmgCarvi fluorescence in starved Mtb-rmgCarvi_int_ (Erdman) populations, averaged over a period of five hours, indicates a clearly bimodal distribution. Red horizontal line represents the median and the dotted red line indicates the quartiles. N=150. (**C**) Ratiometric rmgCarvi fluorescence (cAMP content) during starvation in Mtb-rmgCarvi_int_ is correlated to the magnitude of cAMP induction (defined as the highest cpGFP/mCherry ratio in the 10 hours following exposure to fresh media), following recovery from starvation. The red line represents the curve fit using the four-parameter logistic regression (variable slope) in GraphPad Prism (R^2^ = 0.97). N=125. (**D**) Ratiometric rmgCarvi fluorescence (cAMP content) during starvation in the cells originating from the “low” cAMP population is inversely correlated to the lag time before re-growth begins (N=22 cells). Pearson correlation coefficient (*r*) for the data is-0.77, *p* < 0.0001. The red line represents linear fit to the data. (**E, F**) Correlation plots of ratiometric rmgCarvi fluorescence (cAMP content) during regrowth from starvation and growth rate in “low” cAMP cells from two independent biological replicates, black circles (N=13 cells, Pearson correlation coefficient, *r* = 0.84, *p* = 0.0003) and white circles (N=18 cells, Pearson correlation coefficient, *r* = 0.67, *p* = 0.0022) (**E**) and in “high” cAMP cells from two independent biological replicates, black circles (N=30 cells, Pearson correlation coefficient, *r* = 0.43, *p* = 0.017) and white circles (N=83 cells, Pearson correlation coefficient, *r* = 0.62, *p* < 0.0001) (**F**).

### Use of rmgCarvi to monitor cAMP dynamics in *M. tuberculosis* during infection and as a tool for drug discovery

Next, we aimed to measure cAMP levels in Mtb during intracellular growth in macrophages. Previous reports have shown that Mtb cAMP levels can increase upon infection of macrophages when compared to Mtb incubated in cell culture media alone (21). Based on these results, Mtb-rmgCarvi_epi_ cultured in 7H9+OADC media was washed and resuspended in standard cell culture media (DMEM-based) and then used to infect RAW264.7 murine macrophages. Quantification of cpGFP:mCherry ratios clearly showed a significant increase in cAMP content upon exposure to cell culture media which further increased upon infection (**Fig. 6A**). This induction was, however, not sustained and the cAMP levels dropped after 3 h and was stably maintained at a lower level (**Fig. 6B**), as measured by flow cytometry. Infection of IFN-γ activated macrophages resulted in increased rmgCarvi fluorescence ratios initially at the 0h time point, but these values declined over time and ultimately reached levels comparable to those in unactivated macrophages (**Fig. 6B**). Because acidic pH has previously been shown to induce cAMP in Mtb, we asked whether phagosomal acidification might be responsible for this early increase in reporter signal. To test this, we treated macrophages with established inhibitors of phagosomal acidification, ammonium chloride (NH_4_Cl) (46) or chloroquine (47), however, this did not significantly alter rmgCarvi ratios (**Fig. 6B, S11A**). Likewise, pre-induction of bacterial cAMP by exposing Mtb-rmgCarvi_epi_ to acidic pH 5.5 prior to infection failed to prevent the increase in rmgCarvi fluorescence in intracellular Mtb (**Fig. S11A**). We also carried out imaging of infected macrophages to capture cell-to-cell heterogeneity in Mtb-rmgCarvi_epi_ fluorescence during infection. RAW264.7 murine macrophages were infected with Mtb-rmgCarvi_epi_ (spinfection and incubation for 1h), following which cells were washed to remove extracellular bacteria and cells imaged at 0, 1, 2, 3, 4 and 24 h timepoints (**Fig. 6C, Fig. S12A**). We detected a significant drop in cAMP levels in Mtb at the 4 h time point. In this experiment, we also analyzed rmgCarvi fluorescence in extracellular Mtb in the vicinity of macrophages. However, no significant drop in cAMP levels was detected in this extracellular subpopulation (**Fig. S11B**). Interestingly, this drop in cAMP levels at 4 hours, was observed only in case of RAW264.7 murine macrophages. Infection of human U937 lung macrophage cell line did not result in any drop in Mtb cAMP levels and in fact we observed a small increase in cAMP levels at the 4-hour time point (**Fig. 6D, Fig. S12B**) in intracellular Mtb and no changes in rmgCarvi fluorescence in the uninfected extracellular subpopulation (**Fig. S11C**).

**Figure 6.**
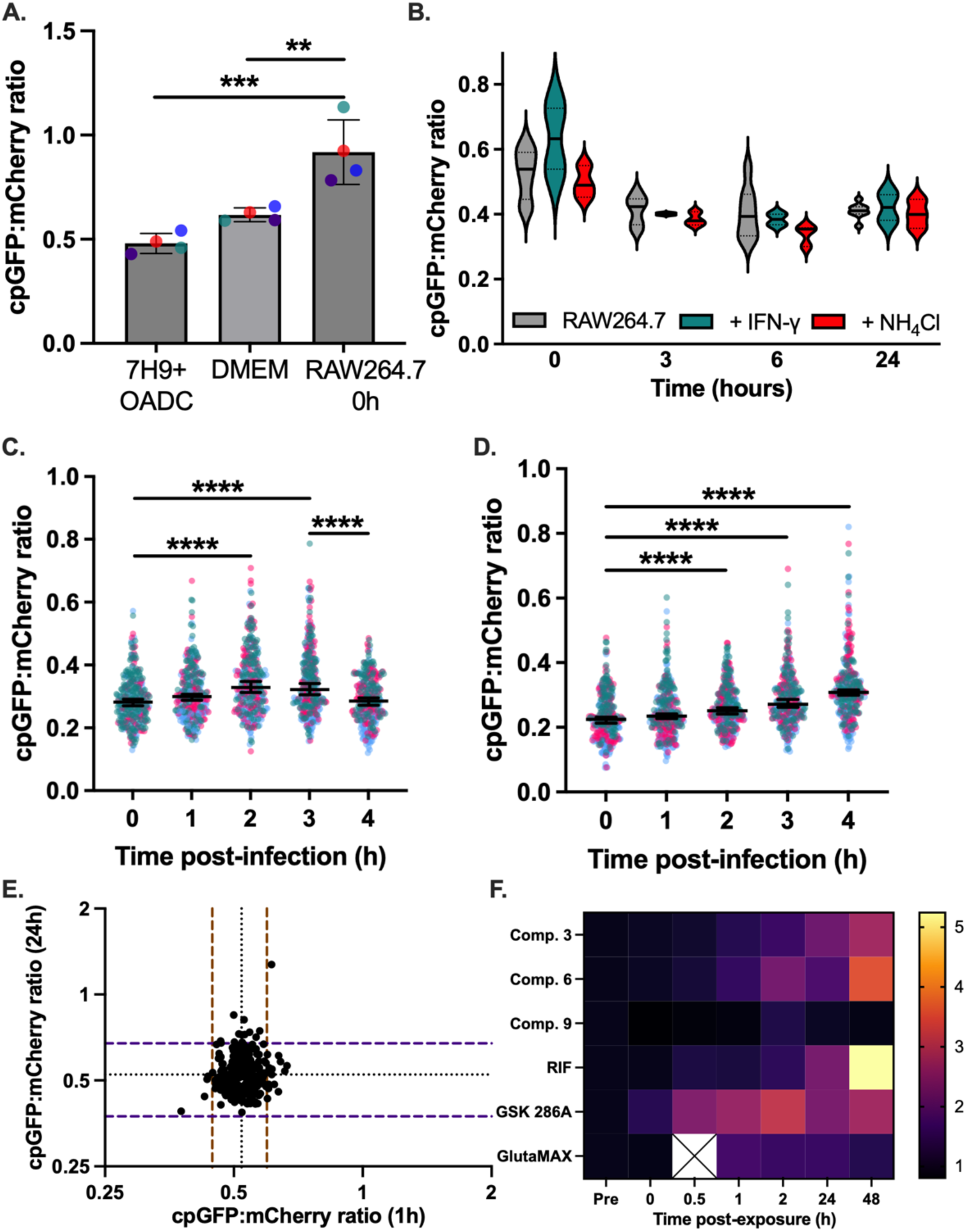
Intra-mycobacterial cAMP dynamics during macrophage infection using Mtb-rmgCarvi. (**A**) Ratiometric rmgCarvi fluorescence of Mtb-rmgCarvi_epi_ (Erdman) cultured in 7H9+OADC or resuspended in DMEM cell culture-based media or immediately upon infection of RAW264.7 macrophages, were measured using a plate reader. cAMP content increases slightly in cell culture media but is significantly increased upon infection (N=4; biological replicates color-coded). Symbols represent means ± SD. One-way ANOVA with Holm-Sidak’s correction ** - *p* < 0.01, *** - *p* < 0.001. (**B**) rmgCarvi fluorescence dynamics in Mtb-rmgCarvi_epi_ during the first 24 hours of infection of RAW264.7 macrophages, with or without IFN-γ activation, or in macrophages where the phagosomal pH was deacidified using 10 mM ammonium chloride (NH_4_Cl), measured by flow cytometry. Intra-mycobacterial cAMP levels drop within 3 hours and is maintained stably over the next 21 hours. cAMP induction upon infection was transient, as indicated by cpGFP and mCherry fluorescence measurements using flow cytometry (N=2-8, 10000 cells per replicate per time point). (**C, D**) Ratiometric rmgCarvi fluorescence values in intracellular Mtb-rmgCarvi_epi_ bacteria upon infection of murine RAW264.7 (**C**) or human U937 (**D**) macrophages, captured by fluorescence microscopy and quantified using FIJI (N=3, >300 events per replicate per time point). Kruskal-Wallis test with Dunn’s correction **** - *p* < 0.0001. (**E**) Mtb-rmgCarvi_epi_, cultured in 96-well plates, were exposed to a library of ∼240 compounds over 2 days. Correlation plot showing the rmgCarvi fluorescence ratios after 1 h vs 24 h exposure. The dotted black horizontal and vertical lines represent the medians at the corresponding time points. The colored lines indicate ±3 median absolute deviation values. (**F**) Heat-map showing the validation of the activity of some of the chosen compounds from the screen. Compounds 3 and 6 were identified as putative cAMP agonists, similar to GSK286 and Rifampicin (RIF). Compound 9 was included as a negative control and GlutaMAX represents the media control. The time points are indicated on the *x-axis*, including the fluorescence prior to exposure to compounds (Pre). All data shown are mean of at least three biological replicates. Biological replicates in panels (**C, D**) are color coded.

Finally, we wanted to evaluate the utility of rmgCarvi to identify small molecules perturbing cAMP homeostasis in Mtb. cAMP-related pathways have recently emerged as potential drug targets in Mtb. Disrupting cAMP signaling restricted bacterial growth in macrophages and impaired pathogenesis by blocking Mtb’s utilization of host lipids (33–35, 48). As a proof of concept, we carried out a small-scale screening of ∼240 prioritized hits from the Global Health Box against Mtb-rmgCarvi_epi_. Measurement of ratiometric rmgCarvi fluorescence allowed us to identify potent molecules that significantly perturbed cAMP levels (**Fig. 6E**). Two hits exhibiting cAMP agonist activity (Comp. 3 and Comp. 6) were further validated in a secondary screen alongside the anti-TB antibiotic rifampicin, the known adenylyl cyclase agonist, GSK286, and a non-hit control (Comp. 9) (**Fig. 6F**).

## Discussion

In this study, we describe the characterization of rmgCarvi as a first-in-class, genetically encoded ratiometric cAMP biosensor, repurposed from a eukaryotic reporter for use in mycobacteria. To our knowledge, this is the first study to characterize cAMP dynamics at the single-cell level in mycobacteria. This biosensor represents a major technical advance that allows for real-time single-cell monitoring of cAMP dynamics. Earlier studies, utilizing bulk-level assays, have established the crucial multi-faceted role of cAMP in Mtb physiology and pathogenesis (6–13). However, these methods failed to capture the cellular heterogeneity and temporal dynamics revealed by rmgCarvi. Our findings establish that intracellular cAMP functions in mycobacteria not just as a static secondary messenger but as a highly tunable metabolite coordinating various physiological processes from carbon utilization, growth rate, cell division and host-cell adaptation.

The use of rmgCarvi in Msm and Mtb revealed species-specific differences in cAMP dynamics. Although Msm showed higher baseline cAMP concentrations, those levels were largely unresponsive to pH, salt or cholesterol, likely reflective of its saprophytic lifestyle vs Mtb’s obligate pathogenic lifestyle. An important insight provided by rmgCarvi is the existence of an “Goldilocks zone” for cAMP, a specific physiological range required for metabolic potency and optimal growth. Our single-cell imaging revealed a direct correlation between cAMP content and growth rate. This observation is consistent with earlier studies which have shown that the cAMP responsive protein (CRP, Rv3676) in Mtb is a global regulator, required for the expression of genes involved in central metabolism, such as the succinate dehydrogenase complex (49) and amino acid biosynthesis (50, 51). Without sufficient cAMP to activate these pathways, Mtb exhibits impaired fitness and reduced growth rates. However, this direct correlation between cAMP content and growth rate cannot be generalized across all growth conditions or environmental conditions. For example, while Mtb bacteria in stationary phase were generally associated with very low cAMP levels, our single-cell analysis revealed significant subpopulations of Mtb with high cAMP both during stationary phase and in Mtb that were starved of nutrients for 6-8 weeks. In addition, high cAMP levels were observed to be growth-prohibitive, particularly in environments with host-relevant carbon sources (cholesterol and fatty acids), consistent with previous reports (8, 20, 33–35). While recovery of Mtb from starvation was accompanied by marked induction in cAMP levels, these spikes were transient, which subsided before growth commenced. The requirement for cAMP to stay within this “Goldilocks zone” likely explains why the loss of the essential adenylyl cyclase, Rv3645 (leading to too little cAMP) and the over-activation of the adenylyl cyclase, Rv1625c (leading to too much cAMP) both results in metabolic dysregulation and growth arrest (8, 33). Reduced cAMP levels in Rv3645 mutants lead to overactive fatty acid import via the Mce1 transporter, resulting in accumulation of toxic intermediates and metabolic derangement (8). Conversely, high cAMP levels are required to activate Mt-Pat (Rv0988), which acetylates and inactivates enzymes like acetyl-Co-A synthetase (20). This acylation prevents the toxic buildup of propionyl-Co-A, a high energy but potentially lethal intermediate of odd-chain fatty acid and cholesterol catabolism. Our observation that rmgCarvi reports differently depending on the carbon source (glucose vs fatty acids) underscores its sensitivity to these metabolic flux changes. Thus, cAMP signaling modulates mycobacterial physiology in a context-dependent manner and by governing central carbon metabolism, functions as a rheostat, that dynamically balances fatty acid uptake and detoxification pathways to support Mtb survival within the host.

The discovery of the transient spike in cAMP preceding cell division in Mtb adds a novel dimension to mycobacterial signaling. However, a key limitation of this observation is defining precisely the timing of cell division, which remains technically challenging without the use of fluorescent septation markers. Nevertheless, the cAMP pulse could function as a potential trigger for cytokinesis, activating downstream effectors involved in cell wall remodeling or septation. In Mtb, the serine/threonine kinase, PknA, phosphorylates several proteins that coordinate cell pole elongation and septal-ring assembly, including the cAMP PDE, Rv0805 (52). PknA-mediated phosphorylation inhibits Rv0805’s enzymatic activity (53) and is also required for the localization of Rv0805 to the mycobacterial cell wall (54). Temporarily inhibiting Rv0805 during this stage could therefore potentially lead to transient increases in intracellular cAMP that may appear as pulses prior to septation, although targeted studies are needed to confirm this connection. Given that cAMP-responsive pathways in Mtb regulate peptidoglycan synthesis and cell wall biogenesis (*mma3, accD3*, *rpfA*) (55), this secondary messenger may directly coordinate cell wall remodeling during septation. Additionally, cAMP could activate protein lysine acetyltransferases like Mt-Pat (20), which could facilitate post-translational regulation of metabolic flux to support the energetic demands of cell division. These observations align with findings in other organisms. Synchronized cultures of the archaeon, *Halobacterium salinarium*, exhibited sharp transient cAMP increases immediately before and after cell division, suggesting a direct role in cell-cycle progression (56). Similarly, synchronized cultures of *E. coli* showed two cAMP peaks per generation: a major peak in mid-cycle coinciding with initiation of DNA synthesis and a smaller peak immediately before or during cell division, suggested to relate to regulation of cell wall formation (57). These data collectively indicate that cAMP oscillations are coordinated with cell-cycle progression across diverse organisms, including peaks around division.

In various bacterial species, the cAMP-CRP regulatory axis functions as a global metabolic signal that coordinates cell division, cell wall synthesis, and cell morphology, linking nutrient status to cell size and shape (58–60). Recent work by Wehrens et al., demonstrated that cAMP-CRP acts as a key feedback mechanism sensing metabolite-dependent signals and dynamically adjusting enzyme expression in real time, suggesting that cAMP may have evolved not only to handle changing external carbon sources but also to buffer or regulate internal metabolic stochasticity (60). While cAMP’s role as a dedicated “cell-cycle timer” is less established than that of c-di-GMP or c-di-AMP (61, 62), its functions as a global metabolic signal influencing division and morphology are well documented in bacteria.

Perhaps the most striking observation from the use of rmgCarvi in Mtb was the detection of bimodal cAMP levels in starved populations, which dictates a distinct hysteresis upon nutrient supplementation. We found that low-cAMP subpopulations experience a prolonged lag time and exhibit strong induction of cAMP, whose magnitude is inversely correlated to cAMP levels during starvation, whereas high-cAMP subpopulations resume growth more rapidly, on average, and exhibit non-correlated induction of cAMP. This establishes cAMP as a metabolic primer for resuscitation. This single-cell evidence provides a mechanistic basis for population-level studies showing that unsaturated fatty acids stimulate resuscitation by activating the adenylyl cyclase, Rv2212, to increase intracellular cAMP (10). Also, cAMP, through CRP, has been shown to activate the transcription of *rpfA*, which encodes the most potent of the five resuscitation-promoting factors in Mtb (44, 63). Rpfs are secreted growth factors that stimulate the recovery of dormant or non-culturable cells, likely by remodeling the peptidoglycan (43). Our results would therefore suggest that Mtb maintains a subpopulation of high-cAMP cells during starvation as a “bet-hedging” strategy to ensure rapid recovery and reactivation upon encountering favorable growth conditions. Interestingly, increased heterogeneity in cAMP levels was also observed in Mtb cultures during stationary phase (**Fig. 3D-F, Table S2**), when nutrients become limiting. The ability to monitor this bimodality in real time reveals how Mtb manages the transition between dormancy and active replication.

The biosensor also successfully captured host-specific cAMP dynamics within macrophages. Strong induction of bacterial cAMP levels was observed upon internalization in both human and murine macrophages; however, only in case of murine RAW macrophages the intracellular Mtb exhibited a drop in cAMP levels 3-4 h post-uptake (**Fig. 6C, D**). The initial cAMP induction and subsequent decline is consistent with the established model of cAMP intoxication, mediated via the adenylyl cyclase Rv0386 (9). This secretion elevates host cAMP levels to subvert innate immunity, specifically by activating host PKA-CREB pathway to inhibit the production of inflammatory cytokines like TNF-α (64, 65). Elevated host cAMP also interferes with phagosome-lysosome fusion by inhibiting actin assembly around the phagosome (22), thereby creating a protected niche for Mtb replication. The subsequent decrease in mycobacterial cAMP seen in murine macrophages may reflect a transition to a persistent state, where the bacteria lower their cAMP as they adapt to the stressful, acidic environment of the phagosome (66). Alternately, the lowered cAMP content might be a requirement consistent with Mtb’s reliance on fatty acids and cholesterol as carbon sources in the intra-phagosomal niche (67, 68). This highlights the biosensor’s utility in deciphering how Mtb tunes its signaling to match the specific microenvironmental cues of the host. Finally, we have demonstrated that rmgCarvi is a robust platform for high-throughput screening of compounds that perturb cAMP levels. By identifying molecules that push Mtb out of its “Goldilocks zone”, either by activating adenylyl cyclases or inhibiting PDEs, we can arrest the pathogen’s growth and enhance its sensitivity to existing antibiotics. The identification and validation of GSK286 and V-59 as promising candidates illustrates the potential for cAMP-targeted therapies to shorten TB treatment durations and overcome intrinsic drug resistance (34, 35, 48).

During this study, we also encountered some limitations with use of rmgCarvi. Although expression of rmgCarvi from either the episomal or integrative constructs did not affect mycobacterial growth and division, the Mtb-rmgCarvi_epi_ strain was not suitable for long-term timelapse imaging. This was likely due to phototoxicity associated with high frequency imaging of two fluorophores (cpGFP and mCherry) in a strain with 3-5x higher reporter expression. Consequently, for all our time-lapse experiments we used the integrative Mtb-rmgCarvi_int_ construct, which although not as bright as the episomal construct emitted fluorescence above the background. Also, while ratiometric rmgCarvi enabled versatile measurement of cAMP across multiple techniques - microscopy, flow cytometry or fluorescent spectrometry, the absolute values were not comparable between methods due to differences in sensitivity and their dynamic ranges.

In conclusion, rmgCarvi provides the necessary cellular and temporal resolution to transform our understanding of how Mtb utilizes this secondary messenger to navigate the complex metabolic and immunological landscape of the human host. Future studies will be aimed at defining the cell cycle-dependent cAMP pulse prior to septum invagination, and the interplay between bacterial and host cAMP signaling during infection. By providing a window into real-time cAMP dynamics in individual bacteria within host cells, rmgCarvi opens the door to mechanistic studies on persistence, immune evasion and metabolic adaptation, ultimately guiding the rational design of interventions that target cAMP-dependent pathways in Mtb. We envision rmgCarvi serving as a template for uncovering conserved, metabolite-regulatory principles in diverse bacterial pathogens with strong implications for antimicrobial development.

## Materials and Methods

### Bacterial Strains and Growth Conditions

*Mycobacterium smegmatis* mc^2^155 (Msm), *Mycobacterium tuberculosis* strains - Erdman, H37Rv, and auxotrophic strain mc^2^6206 (Mtb), were grown in Middlebrook 7H9 broth base media (Difco) at 37°C in an orbital shaker set at 100 rpm. Where indicated, the 7H9 broth base media was supplemented with 10% ADS (final concentrations: 0.5% bovine albumin fraction V, 0.2% dextrose, 0.085% sodium chloride, all from Sigma), 10% OADC (0.5% bovine albumin fraction V, 0.2% dextrose, 0.0004% catalase, 0.005% oleic acid, and 0.085% sodium chloride, BD BBL), 10% AN-BCP (BCP media: 0.5% fatty acid-free bovine albumin fraction V, 0.085% sodium chloride, 0.1% w/v butyric acid, 0.01% w/v cholesterol, and 0.001% w/v palmitic acid). Where indicated, various other carbon sources were supplemented to the 7H9 broth base, including glycerol (0.5%), dextrose (0.2%), oleic acid (0.2 mM), and propionate (100µM). All Mtb mc^2^6206 cultures were additionally supplemented with 24 µg/mL L-pantothenate and 50 µg/mL L-leucine. All broth cultures were supplemented with 0.02% tyloxapol. When necessary, CRISPRi knockdown strains were induced with 100 ng/mL anhydrotetracycline (aTc; Takara Bio) for two days before assay. For culturing of bacteria on solid media or when generating recombinant strains from parental Msm, or Mtb strains, Middlebrook 7H10 agar (Difco Middlebrook 7H10 powder base, 0.5% glycerol, and 10% OADC supplement) containing either 15 µg /mL kanamycin, 50 µg /mL hygromycin B, or both, was used. After incubating the plates at 37°C (3 days in case of Msm and 3 weeks in case of Mtb), single colonies were picked, grown to mid-exponential phase in 7H9 + 10% ADS media, and aliquots prepared that were kept frozen at - 80°C. Individual aliquots were used once or twice and discarded.

### Construction of reporter strains

Primers, plasmids, and strains made and/or used in this study are listed in **Supplementary Table 1**. pUC57-rm_gCarvi-T2A was a gift from Naoto Saitoh (Addgene plasmid # 190334; http://n2t.net/addgene:190334; RRID:Addgene_190334). All cloning procedures were carried out in TOP10 *E. coli* grown in standard LB media containing the appropriate antibiotics (kanamycin - 50 µg /mL; hygromycin - 150 µg /mL; ampicillin - 100 µg /mL). The gCarvi fragment was PCR amplified using the rmgCarvi-hsp60-NheI-F and gCarvi-ScaI-R primers. These PCR products carrying overhangs encoding the relevant restriction sites for cloning were cloned into the pCR2.1-TOPO vector (Invitrogen). Plasmid isolation, PCR and/or gel purification was carried out using Qiagen miniprep kits, QIAquick PCR and gel clean-up kit, respectively, as per manufacturer’s instructions (Qiagen). Upon confirmation of the rmgCarvi sequence by Sanger sequencing, the fragment was sub-cloned into the pND257 plasmid (69) by digesting both pCR2.1-TOPO-rmgCarvi and pND257 with the NheI and ScaI restriction enzymes, followed by T4 DNA ligation, transformation into TOP10 *E. coli*, colony selection, plasmid purification, and whole plasmid sequence confirmation. Plasmid DNA was then electroporated into either *M. smegmatis* mc^2^155, *M. tuberculosis* mc^2^6206, *M. tuberculosis* strain H37Rv, or *M. tuberculosis* strain Erdman. The pMV261-rmgCarvi plasmid was constructed by digesting pMV261(70) with the MluI and MscI restriction enzymes and pND257-rmgCarvi with the MluI and ScaI restriction enzymes, followed by gel purification of the relevant bands. The rmgCarvi and pMV261 plasmid DNA fragments were then ligated using T4 DNA ligase, transformed into TOP10 *E. coli* cells, and confirmed by restriction digest mapping and sequencing the whole plasmid construct. pMV261-rmgCarvi was electroporated into either *M. smegmatis* mc^2^155, *M. tuberculosis* mc^2^6206, *M. tuberculosis* strain H37Rv, or *M. tuberculosis* strain Erdman. Site-directed mutagenesis of the rmgCarvi gene was carried out using a Q5 Site-directed mutagenesis kit (New England Biolabs) according to manufacturer instructions. After verification of the amplified fragments by sequencing, the fragments were excised using the appropriate restriction enzymes to sub-clone into the pMV261 plasmids. Correct recombinant clones, verified either by restriction digestion or sequencing, were electroporated into Msm or Mtb (Bio-Rad GenePulser Xcell; 2.5 kV, 1000 Ohms, 25 µFD) and plated on Middlebrook 7H10 agar as described above.

### Macrophage Infection

Propagation and infection of RAW264.7 (ATCC: TIB-71) and U937 cells (ATCC: CRL-1593.2) was done as previously described, with some modifications (71). Laboratory stock aliquots of RAW264.7 murine macrophage and U937 human monocytes were thawed and sub-cultured in Dulbecco’s modified Eagle medium (DMEM; Invitrogen), and Roswell Park Memorial Institute 1640 medium (RPMI; Invitrogen), respectively, supplemented with 10% fetal bovine serum (FBS; Invitrogen) for three days in a tissue culture-treated T-75 flask (Corning). Cells were then detached, pelleted, re-suspended in 5 mL fresh, pre-warmed growth media. Cell counts and viability assessment was carried out using Trypan blue staining (0.4%; Gibco). About 1.2 x 10^5^ viable U937 cells in a volume of 300 µL were seeded into each well of an 8-chambered ibidi µ-dish (ibidi) and were treated with phorbol 12-myristate-13-acetate (PMA) at a final concentration of 50 ng/mL, 48 hours prior to infection. After 24 hours of differentiation into macrophages, the cells were washed with phosphate-buffered saline (PBS; Invitrogen) three times, and the cells allowed to rest for 24 hours in fresh RPMI 1640 media lacking PMA. About 0.8 x 10^5^ viable RAW264.7 macrophages were seeded into each well of an 8-chambered Ibidi µ-dish (Ibidi), 24 hours prior to infection for cells to adhere. The following day, 10 mL of a mid-exponential (A_600 nm_ 0.3-0.6) Mtb-rmgCarvi_epi_ culture was pelleted by centrifugation at 5000 x g for five minutes. Bacterial pellets were resuspended in 1 mL fresh phenol-red free DMEM or RPMI 1640 media, supplemented with 10% FBS, 10mM HEPES, 1mM sodium pyruvate, and GlutaMAX (all Invitrogen).

For infections analyzed by wide-field fluorescence microscopy (Leica Microsystems), the bacterial samples were passed through a 5 µm filter to produce a single-cell suspension. The single cell bacterial suspensions were added to the macrophages in a minimal volume of 130 µL per well of the 8-chambered ibidi µ-dish. The infection mix was subject to low-speed centrifugation (spinfection; 500 x g, 5 min) to improve the uptake of the bacteria by macrophages. Following one hour of incubation at 37°C and 5% CO_2_, the extracellular bacteria were removed by washing three times with pre-warmed PBS; representing the 0 h timepoint. The infection was imaged using a Leica Thunder Imaging system equipped with a scientific CMOS Leica K8 camera and an HC PL Fluotar 100X/1.32 oil immersion objective (Leica). Images were acquired every hour till 6h post-infection, and finally at the 24h post-infection timepoint.

In case of flow cytometry, the bacterial samples were subjected to low-speed centrifugation (80 x g, 5 min) to pellet large clumps and generate a single-cell suspension. The single-cell suspension was diluted to infect RAW264.7 cells at an MOI of 10. The bacteria were allowed to adhere to macrophages for 3.5 hours after which the non-adherent bacteria were removed by washing three times with pre-warmed PBS and cultured in fresh media; representing the 0h timepoint. The macrophages were collected and subjected to flow cytometry measurements (see below) at the indicated timepoints, using a BD FACS Aria Fusion flow cytometer. At least 10,000 events, for each of 4 technical replicates, were analyzed per timepoint per biological replicate.

### Characterization of rmgCarvi in mycobacteria

Wild-type Msm, Msm-rmgCarvi_epi_ and Msm-rmgCarvi_int_ strains were cultured to mid-exponential phase (A_600 nm_ 0.3-0.6) in 7 mL standard 7H9 + 10% ADS axenic media. About 2 OD_600_ equivalent of cells were pelleted and re-suspended in 1 ml lysis buffer (50 mM Tris-HCl, 150 mM NaCl, 1 mM EDTA, 10% glycerol, and 0.05% Triton X-100) and mechanically disrupted by bead-beating, 5 x 1 min, with zirconia beads (0.1 mm; BioSpec Products). Lysates were centrifuged at 15000 x g for 5 minutes to pellet cell debris, and 100 µl of the supernatant was aliquoted to individual wells of a 96-well plate. Bacterial lysates were exposed to varying concentrations (0.01-1000 μM) of adenosine 3’-5’ cyclic monophosphate (cyclic AMP; Sigma) in total volume of 200 µl per well. The plate was incubated for 30 minutes at 37°C, and fluorescence readings (cpGFP:Ex_500 nm_, Em_525 nm_; mCherry:Ex_587 nm_, Em_612 nm_) were taken on a SpectraMax i3x plate reader (Molecular Devices). Fold-change and dynamic ranges were obtained by first subtracting fluorescence values obtained from wild-type Msm strain and normalizing to minimum fluorescence obtained at the lowest concentration of cAMP (*F_min_*). Each strain was analyzed using three independent biological replicates.

### Generation and validation of individual CRISPRi strains

sgRNAs targeting the respective ORF were selected using the previously published sgRNA design tool for mycobacteria (https://pebble.rockefeller.edu/tools/sgrna-design/)(72). Two complementary oligonucleotides with BsmBI overhangs were ordered (IDT), annealed and cloned into BsmBI-digested, phage L5-integrative, pIRL2 plasmid (plRL2 was a gift from Jeremy Rock (Addgene plasmid # 163631; http://n2t.net/addgene:163631; RRID:Addgene_163631). Individual CRISPRi plasmids, after verification by sequencing, were electroporated into Mtb-rmgCarvi_epi,_ as described before. Total RNA was extracted from Mtb-rmgCarvi_epi_ transformants harboring the individual Mtb CRISPRi constructs, to estimate the efficiency of genetic knockdown, by qPCR (72). Briefly, early exponential Mtb-rmgCarvi_epi_ + pIRL2-CRISPRi cultures were left uninduced or induced with 100 ng/mL aTc for two days. After the induction period, bacterial cultures were pelleted, washed once in 1X PBS, and resuspended in 500 µl TRIzol (Invitrogen). Cell lysis was achieved by bead beating (5 x 1 min) using 250 µl zirconia beads, in a bead beater (Biospec). The samples were centrifuged to pellet the beads and cell debris, and the supernatant was extracted with 50 µl 1-bromo-3-chloro propane. Total RNA was precipitated from the aqueous phase with 250 µl isopropanol and the precipitated RNA was pelleted and washed with 70% ethanol twice before re-suspension in nuclease free water (Ambion). RNA quality and concentrations were measured using a NanoDrop One UV/Vis spectrophotometer, and ∼500 ng of total RNA was treated with DNase I using the TURBO DNA-free kit (Invitrogen), according to manufacturer’s protocol. cDNA synthesis was carried out using the SuperScript IV first-strand synthesis system (Invitrogen). Approximately 50 ng of the resulting cDNA was used for quantitative real time polymerase chain reaction (qRT-PCR) using iQ SYBR Green Supermix (Bio-Rad) and the CFX Connect Real-Time qRT-PR System (Bio-Rad).

### Immunoblotting

Bacteria from exponential and stationary phase cultures of Mtb mc^2^6206, and the transformants Mtb-rmgCarvi_int_ and Mtb-rmgCarvi_epi_ were harvested and protein lysates prepared using bead beating as described above for Msm. Protein concentrations were quantified using the bicinchoninic (BCA) assay (Pierce; Thermo Scientific). Protein samples (15 μg) were separated on Novex pre-cast 4-12% Tris-glycine gels (Thermo Scientific) and transferred to nitrocellulose membranes (Thermo Scientific) via semi-dry transfer using a Trans-blot SD semi-dry transfer cell (Bio-Rad) according to manufacturer instructions. Membranes were briefly rinsed in 1X TBST (20 mM Tris, 150 mM NaCl, and 0.1% Tween-20, pH 7.6) and blocked for 1 hour in Intercept blocking buffer (LICORbio). Monoclonal anti-FLAG M2 (Sigma-Aldrich) was diluted 1:1,000 in blocking buffer + 0.5X TBST and incubated with membranes overnight at 4°C. Following five washes (5 minutes each) in 1X TBST buffer, membranes were incubated with secondary antibody (source: IRDye 680RD goat anti-mouse, diluted 1:10,000) for one hour at room temperature. Membranes were washed 5 times (5 minutes each) in 1X TBST and imaged using an Odyssey CLx Imaging System (LICORbio).

### Fluorescence measurements using spectrometry

All plate reader assays were carried out in Costar 3603 96-well assay plates (Corning) using a SpectraMax i3x plate reader and SoftMax Pro 7.2 software (Molecular Devices). About 200 µl of bacterial cultures (Msm and Mtb wild type and rmgCarvi expressing strains) were aliquoted into individual wells in triplicate, avoiding the border wells. cpGFP fluorescence was measured at Ex_500 nm_ and Em_525 nm_ (Bandwidth: excitation = 9 nm, emission = 15 nm). mCherry fluorescence was measured at Ex_587 nm_ and Em_612 nm_ (Bandwidth: excitation = 9 nm, emission = 15 nm). All reads were performed using the following parameters: PMT gain set to “High”, 6 flashes per read and a read distance of 1 mm from the plate bottom. Background fluorescence was measured using A_600 nm_ matched wild-type Msm/Mtb cultures to enable OD-normalized background subtraction.

For determination of fluorescence spectra, protein lysates from Mtb-rmgCarvi_epi_, prepared as described above, were aliquoted into Costar 3603 96 well assay plates (Corning). Emission spectra were recorded between 525 nm and 580 nm, with 5 nm increments, for cpGFP (Ex_500 nm_) and between 612 nm and 747 nm, with 5 nm increments, for mCherry (Ex_587 nm_) and excitation spectra between 350 nm and 500 nm, with 5 nm increments, for cpGFP (Em_525 nm_) and between 435 nm and 585 nm, with 5 nm increments, for mCherry (Em_612 nm_),

### Determination of cAMP levels by ELISA

cAMP levels were measured using an ELISA-based method as described previously, with some modifications (8). Briefly, mid-exponential cultures of Mtb-rmgCarvi_epi_ were left untreated or exposed to the cAMP agonist, GSK286 (5 µM), for four hours. In case of CRISPRi strains, Mtb cultures were induced with 100 ng/mL aTc for two days, prior to the assay. Protein lysates were prepared as described above for Msm, and protein concentration measured by BCA assay (Pierce; Thermo Scientific). cAMP content, normalized to total protein, was measured using a Cyclic AMP ELISA Kit (Caymen Chemicals) according to manufacturer instructions. Each strain was analyzed using three independent biological replicates.

### Flow Cytometry

Flow cytometry was performed using a BD FACS Aria Fusion flow cytometer equipped with 488nm and 561nm lasers. The FITC (530/30-nm) and PE-Texas Red-mCherry-PI (610/20-nm) emission channels were used to detect the emission wavelengths from the 488 nm and 561 nm lasers, respectively. The cells were first gated on the forward versus side scatter plot to identify bacteria and to exclude debris and then gated for singlet cells by gating along the main diagonal. The singlet population was then analyzed for their FITC and PE fluorescence in their respective plots. The fluorescence thresholds were gated using the non-fluorescent parental wildtype Mtb H37Rv strain, and single fluorescent GFP-expressing and tdTomato-expressing Mtb H37Rv strains, as positive controls for fluorescence. The ratio of the median FITC fluorescence to the median PE fluorescence in the subpopulation of singlet cells positive for PE fluorescence (indicating that these cells express the mCherry fluorophore and are, therefore, rmgCarvi expressing cells) was used as a measure of the intracellular cAMP concentration. At least 10,000 events per replicate were analyzed for all experiments.

### Microscopy

In case of axenic cultures, at each time point during the different growth phases, images were captured using a Leica Thunder Imaging system equipped with a scientific CMOS Leica K8 camera and an HC PL Fluotar 100X/1.32 oil immersion objective. Cells were imaged within 1-2 min after spotting them on agarose pads. In addition to phase images, cpGFP fluorescence was captured using Ex_475 nm_/Em_535 nm_, 25% power, 100 ms exposure; and mCherry fluorescence was captured using Ex_575 nm_/Em_642 nm_, 15% power, 100 ms exposure. In case of infected cells (RAW264.7 and U937 macrophages), exposure times for cpGFP and mcherry channels were 50 ms and 25 ms, respectively. Images were acquired using LAS-X acquisition software (Leica) and processed using FIJI (73). For each condition, at least three independent biological replicates were sampled and 200–600 cells were imaged.

### Time-lapse Microscopy

Time-lapse microscopy experiments were carried out as described previously, with some modifications (41). Bacteria from exponentially growing cultures of Mtb-rmgCarvi_int_ were seeded into a custom-made microfluidic device as described previously, with the only difference being that we used an agarose-based membrane instead of cellulose membrane (41). Media (7H9+ADS or 7H9+OADC) was perfused through the device using a syringe pump (Aladdin syringe pump, World Precision Instruments), at a flow rate of 10-15 μl/min. Bacteria in about 30-45 independent xy positions were imaged using an HC PL Fluotar 100X/1.32 oil immersion objective at 1 h intervals, over a period of 7 days. When imaging starved Mtb bacteria, the cells were seeded undiluted and imaged for a period of at least 20 h in the starvation medium (PBS + 0.02% tyloxapol) before exposing them to fresh media. Images were captured on phase (100 ms exposure), cpGFP (Ex_475 nm_/Em_535 nm_, 20% power, 50 ms exposure) and mCherry channels (Ex_575 nm_/Em_642 nm_, 15% power, 25 ms exposure) using the LAS-X acquisition software. Acquired images were exported to FIJI for further processing and analysis.

### Image Analysis

Segmentation and analysis of snapshots and image sequences was done using FIJI (73) or CellProfiler (74). In each case, fluorescence was measured by thresholding then masking cells in the mCherry channel, then using the mask to measure fluorescence in the cpGFP channel. Background fluorescence was measured separately, from cell-free regions, for each image and subtracted from the cpGFP/mCherry fluorescence intensities of each cell in that image. Cell length was measured using CellProfiler and the MeasureObjectSizeShape module (74). For time-lapse image analysis, cells across channels and frames were aligned using the MicrobeJ plug-in and the HyperStackRegPlus module (75).

## Screening of compound library

A library of ∼240 compounds with confirmed activity against infections, mosquito-borne diseases and vectors of global concern (Global Health Priority Box) was obtained from Medicines for Malaria Venture (MMV). The compounds were screened at a concentration of 10 µM against Mtb-rmgCarvi_epi_ bacteria seeded in 96-well plates. Compounds causing more than 3 median absolute deviation from the median ratiometric rmgCarvi fluorescence values at 1 h or 24 h timepoints were considered as hits. Retesting of the hit compounds was subsequently carried out (two biological replicates) with more timepoints in secondary assays to confirm activity.

## Statistical Analyses

Most statistical analyses were performed in GraphPad Prism (version 10.6.1 for Windows 11/Mac OS X, GraphPad Software, Boston, Massachusetts USA, www.graphpad.com). For correlations, linear regression analysis was performed with default parameters. Unpaired t tests were performed assuming Gaussian distribution as well as equal standard deviation between samples. Hartigan’s dip test for unimodality was performed using the diptest Python module.

## Acknowledgments

We would like to acknowledge Michelle Gerber for excellent technical support. We thank Drs. William R. Jacobs and Catherine Vilcheze for providing the Mtb mc^2^6206 strain. We acknowledge Medicines for Malaria Venture (MMV) for providing the Global Health Priority Box compound library and GSK (David Barros and Joël Lelievre) for providing the GSK2556286 compound. This research was supported by funds from Canadian Institutes of Health Research (ARB-185715 and ARB-192058); Natural Sciences and Engineering Research Council of Canada (RGPIN-2023-05746) and Saskatchewan Health Research Foundation (6239) to ND. VIDO receives operational funding from the Government of Saskatchewan through Innovation Saskatchewan and the Ministry of Agriculture and from the Canada Foundation for Innovation through the Major Science Initiatives Fund.

## Author Contributions

SW and ND designed research. SW, SSK and ND performed research. AS contributed new reagents or analytic tools. SW, SSK and ND analyzed data and wrote the paper.

## Notes

### Competing Interest Statement

The authors have declared no competing interest.

## References

1. M. Silva, A. J. Pinto, T. Beites, Essential redundancies fuel Mycobacterium tuberculosis adaptation to the host. PLOS Pathogens 21, e1013749 (2025).

2. D. F. Warner, A. K. Barczak, M. G. Gutierrez, V. Mizrahi, Mycobacterium tuberculosis biology, pathogenicity and interaction with the host. Nat Rev Microbiol 1–17 (2025). 10.1038/s41579-025-01201-x.

3. S. S. Kurpad, N. Dhar, Playing Telephone: How Secondary Messengers Influence Host–Pathogen Interactions in Tuberculosis. ACS Infect. Dis. (2025). 10.1021/acsinfecdis.5c00077.

4. R. M. Johnson, K. A. McDonough, Cyclic nucleotide signaling in Mycobacterium tuberculosis: an expanding repertoire. Pathog Dis 76, fty048 (2018).

5. K. A. McDonough, A. Rodriguez, The myriad roles of cyclic AMP in microbial pathogens: from signal to sword. Nat Rev Microbiol 10, 27–38 (2011).

6. G. S. Knapp, K. A. McDonough, Cyclic AMP Signaling in Mycobacteria. Microbiol Spectr 2 (2014).

7. D. Kathayat, B. C. VanderVen, Exploiting cAMP signaling in Mycobacterium tuberculosis for drug discovery. Trends Microbiol S0966–842X(24)00008–8 (2024). 10.1016/j.tim.2024.01.008.

8. A. I. Wong, et al., Cyclic AMP is a critical mediator of intrinsic drug resistance and fatty acid metabolism in M. tuberculosis. eLife 12, e81177 (2023).

9. N. Agarwal, G. Lamichhane, R. Gupta, S. Nolan, W. R. Bishai, Cyclic AMP intoxication of macrophages by a Mycobacterium tuberculosis adenylate cyclase. Nature 460, 98–102 (2009).

10. A. Abdel Motaal, I. Tews, J. E. Schultz, J. U. Linder, Fatty acid regulation of adenylyl cyclase Rv2212 from Mycobacterium tuberculosis H37Rv. FEBS J 273, 4219–4228 (2006).

11. M. O. Shleeva, et al., Overexpression of Adenylyl Cyclase Encoded by the Mycobacterium tuberculosis Rv2212 Gene Confers Improved Fitness, Accelerated Recovery from Dormancy and Enhanced Virulence in Mice. Frontiers in Cellular and Infection Microbiology 7 (2017).

12. M. Thomson, et al., Expression of a novel mycobacterial phosphodiesterase successfully lowers cAMP levels resulting in reduced tolerance to cell wall–targeting antimicrobials. J Biol Chem 298, 102151 (2022).

13. A. R. Shenoy, N. Sreenath, M. Podobnik, M. Kovačevič, S. S. Visweswariah, The Rv0805 Gene from Mycobacterium tuberculosis Encodes a 3‘,5‘-Cyclic Nucleotide Phosphodiesterase: Biochemical and Mutational Analysis. Biochemistry 44, 15695–15704 (2005).

14. B. Görke, J. Stülke, Carbon catabolite repression in bacteria: many ways to make the most out of nutrients. Nat Rev Microbiol 6, 613–624 (2008).

15. A. Kolb, S. Busby, H. Buc, S. Garges, S. Adhya, TRANSCRIPTIONAL REGULATION BY cAMP AND ITS RECEPTOR PROTEIN. Annual Review of Biochemistry 62, 749–797 (1993).

16. H. Khan, et al., Mycobacterium tuberculosis PhoP integrates stress response to intracellular survival by regulating cAMP level. eLife 13, RP92136 (2024).

17. G. Bai, L. A. McCue, K. A. McDonough, Characterization of Mycobacterium tuberculosis Rv3676 (CRPMt), a Cyclic AMP Receptor Protein-Like DNA Binding Protein. Journal of Bacteriology 187, 7795–7804 (2005).

18. C. Kahramanoglou, et al., Genomic mapping of cAMP receptor protein (CRPMt) in Mycobacterium tuberculosis: relation to transcriptional start sites and the role of CRPMt as a transcription factor. Nucleic Acids Research 42, 8320–8329 (2014).

19. M. Stapleton, et al., Mycobacterium tuberculosis cAMP Receptor Protein (Rv3676) Differs from the Escherichia coli Paradigm in Its cAMP Binding and DNA Binding Properties and Transcription Activation Properties. J Biol Chem 285, 7016–7027 (2010).

20. S. Nambi, et al., Cyclic AMP-dependent protein lysine acylation in mycobacteria regulates fatty acid and propionate metabolism. J Biol Chem 288, 14114–14124 (2013).

21. G. Bai, D. D. Schaak, K. A. McDonough, cAMP levels within Mycobacterium tuberculosis and M. bovis BCG increase upon infection of macrophages. FEMS Immunol Med Microbiol 55, 68–73 (2009).

22. S. A. Kalamidas, et al., cAMP synthesis and degradation by phagosomes regulate actin assembly and fusion events: consequences for mycobacteria. J Cell Sci 119, 3686–3694 (2006).

23. Z. Zhang, X. Cheng, Y. Zhao, Y. Yang, Lighting Up Live-Cell and In Vivo Central Carbon Metabolism with Genetically Encoded Fluorescent Sensors. Annual Review of Analytical Chemistry 13, 293–314 (2020).

24. S. Kawata, Y. Mukai, Y. Nishimura, T. Takahashi, N. Saitoh, Green fluorescent cAMP indicator of high speed and specificity suitable for neuronal live-cell imaging. Proceedings of the National Academy of Sciences 119, e2122618119 (2022).

25. K. Harada, et al., Red fluorescent protein-based cAMP indicator applicable to optogenetics and in vivo imaging. Sci Rep 7, 7351 (2017).

26. C. I. Massengill, et al., Sensitive genetically encoded sensors for population and subcellular imaging of cAMP in vivo. Nat Methods 19, 1461–1471 (2022).

27. C. R. Hackley, E. O. Mazzoni, J. Blau, cAMPr: A single-wavelength fluorescent sensor for cyclic AMP. Science Signaling 11, eaah3738 (2018).

28. M. Zaccolo, et al., A genetically encoded, fluorescent indicator for cyclic AMP in living cells. Nat Cell Biol 2, 25–29 (2000).

29. L. M. DiPilato, X. Cheng, J. Zhang, Fluorescent indicators of cAMP and Epac activation reveal differential dynamics of cAMP signaling within discrete subcellular compartments. Proceedings of the National Academy of Sciences 101, 16513–16518 (2004).

30. G. Kudla, A. W. Murray, D. Tollervey, J. B. Plotkin, Coding-Sequence Determinants of Gene Expression in Escherichia coli. Science 324, 255–258 (2009).

31. J. R. McDowell, et al., Mycobacterial phosphodiesterase Rv0805 is a virulence determinant and its cyclic nucleotide hydrolytic activity is required for propionate detoxification. Molecular Microbiology 119, 401–422 (2023).

32. B. C. VanderVen, et al., Novel inhibitors of cholesterol degradation in Mycobacterium tuberculosis reveal how the bacterium’s metabolism is constrained by the intracellular environment. PLoS Pathog 11, e1004679 (2015).

33. R. M. Johnson, et al., Chemical activation of adenylyl cyclase Rv1625c inhibits growth of Mycobacterium tuberculosis on cholesterol and modulates intramacrophage signaling. Mol Microbiol 105, 294–308 (2017).

34. K. M. Wilburn, et al., Pharmacological and genetic activation of cAMP synthesis disrupts cholesterol utilization in Mycobacterium tuberculosis. PLoS Pathog 18, e1009862 (2022).

35. K. L. Brown, et al., Cyclic AMP-Mediated Inhibition of Cholesterol Catabolism in Mycobacterium tuberculosis by the Novel Drug Candidate GSK2556286. Antimicrob Agents Chemother 67, e0129422 (2023).

36. A. M. Gronenborn, R. Sandulache, S. Gärtner, G. M. Clore, Mutations in the cyclic AMP binding site of the cyclic AMP receptor protein of Escherichia coli. Biochem J 253, 801–807 (1988).

37. S. Li, et al., CRISPRi chemical genetics and comparative genomics identify genes mediating drug potency in Mycobacterium tuberculosis. Nat Microbiol 7, 766–779 (2022).

38. S. Rebollo-Ramirez, G. Larrouy-Maumus, NaCl triggers the CRP-dependent increase of cAMP in *Mycobacterium tuberculosis*. Tuberculosis 116, 8–16 (2019).

39. J. Green, et al., Cyclic-AMP and bacterial cyclic-AMP receptor proteins revisited: adaptation for different ecological niches. Curr Opin Microbiol 18, 1–7 (2014).

40. L. P. S. de Carvalho, et al., Metabolomics of *Mycobacterium tuberculosis* Reveals Compartmentalized Co-Catabolism of Carbon Substrates. Chemistry & Biology 17, 1122–1131 (2010).

41. N. Dhar, G. Manina, Single-cell analysis of mycobacteria using microfluidics and time-lapse microscopy. Methods Mol Biol 1285, 241–256 (2015).

42. M. Shleeva, et al., Cyclic Amp-Dependent Resuscitation of Dormant Mycobacteria by Exogenous Free Fatty Acids. PLOS ONE 8, e82914 (2013).

43. M. O. Shleeva, G. R. Demina, A. S. Kaprelyants, Biochemistry of Reactivation of Dormant Mycobacteria. Biochemistry Moscow 90, S193–S213 (2025).

44. L. Rickman, et al., A member of the cAMP receptor protein family of transcription regulators in Mycobacterium tuberculosis is required for virulence in mice and controls transcription of the rpfA gene coding for a resuscitation promoting factor. Mol Microbiol 56, 1274–1286 (2005).

45. J. C. Betts, P. T. Lukey, L. C. Robb, R. A. McAdam, K. Duncan, Evaluation of a nutrient starvation model of Mycobacterium tuberculosis persistence by gene and protein expression profiling. Mol Microbiol 43, 717–731 (2002).

46. P. D. Hart, M. R. Young, Ammonium chloride, an inhibitor of phagosome-lysosome fusion in macrophages, concurrently induces phagosome-endosome fusion, and opens a novel pathway: studies of a pathogenic mycobacterium and a nonpathogenic yeast. Journal of Experimental Medicine 174, 881–889 (1991).

47. M. Mauthe, et al., Chloroquine inhibits autophagic flux by decreasing autophagosome-lysosome fusion. Autophagy 14, 1435–1455 (2018).

48. E. L. Nuermberger, et al., GSK2556286 Is a Novel Antitubercular Drug Candidate Effective In Vivo with the Potential To Shorten Tuberculosis Treatment. Antimicrobial Agents and Chemotherapy 66, e00132–22 (2022).

49. G. S. Knapp, et al., Role of intragenic binding of cAMP responsive protein (CRP) in regulation of the succinate dehydrogenase genes Rv0249c-Rv0247c in TB complex mycobacteria. Nucleic Acids Res 43, 5377–5393 (2015).

50. Y. Liu, S. Rebollo-Ramirez, G. Larrouy-Maumus, Metabolomics reveals that the cAMP receptor protein regulates nitrogen and peptidoglycan synthesis in Mycobacterium tuberculosis. RSC Adv. 10, 26212–26219 (2020).

51. G. Bai, D. D. Schaak, E. A. Smith, K. A. McDonough, Dysregulation of serine biosynthesis contributes to the growth defect of a Mycobacterium tuberculosis crp mutant. Molecular Microbiology 82, 180–198 (2011).

52. M. Thakur, P. K. Chakraborti, GTPase Activity of Mycobacterial FtsZ Is Impaired Due to Its Transphosphorylation by the Eukaryotic-type Ser/Thr Kinase, PknA *. Journal of Biological Chemistry 281, 40107–40113 (2006).

53. N. Malhotra, S. Karthikeyan, P. K. Chakraborti, Phosphorylation of mycobacterial phosphodiesterase by eukaryotic-type Ser/Thr kinase controls its two distinct and mutually exclusive functionalities. Journal of Biological Chemistry 292, 17362–17374 (2017).

54. N. Malhotra, P. K. Chakraborti, Eukaryotic-Type Ser/Thr Protein Kinase Mediated Phosphorylation of Mycobacterial Phosphodiesterase Affects its Localization to the Cell Wall. Front. Microbiol. 7 (2016).

55. J. Barba, A. H. Alvarez, M. A. Flores-Valdez, Modulation of cAMP metabolism in *Mycobacterium tuberculosis* and its effect on host infection. Tuberculosis 90, 208–212 (2010).

56. A. Baumann, C. Lange, J. Soppa, Transcriptome changes and cAMP oscillations in an archaeal cell cycle. BMC Cell Biol 8, 21 (2007).

57. A. V. Lazareva, R. B. Shiian, I. V. Evtodienko, [Changes in cAMP levels in bacterial cells during the cell cycle]. Biokhimiia 52, 1469–1473 (1987).

58. L. Liu, et al., Cyclic AMP-CRP Modulates the Cell Morphology of Klebsiella pneumoniae in High-Glucose Environment. Front. Microbiol. 10 (2020).

59. R. D’Ari, A. Jaffé, P. Bouloc, A. Robin, Cyclic AMP and cell division in Escherichia coli. Journal of Bacteriology 170, 65–70 (1988).

60. M. Wehrens, L. H. J. Krah, B. D. Towbin, R. Hermsen, S. J. Tans, The interplay between metabolic stochasticity and cAMP-CRP regulation in single E. coli cells. Cell Reports 42 (2023).

61. A. Duerig, et al., Second messenger-mediated spatiotemporal control of protein degradation regulates bacterial cell cycle progression. Genes Dev. 23, 93–104 (2009).

62. K. A. Selim, et al., Diurnal metabolic control in cyanobacteria requires perception of second messenger signaling molecule c-di-AMP by the carbon control protein SbtB. Science Advances 7, eabk0568 (2021).

63. G. V. Mukamolova, et al., A family of autocrine growth factors in Mycobacterium tuberculosis. Molecular Microbiology 46, 623–635 (2002).

64. C. M. Leopold Wager, et al., Activation of transcription factor CREB in human macrophages by Mycobacterium tuberculosis promotes bacterial survival, reduces NF-kB nuclear transit and limits phagolysosome fusion by reduced necroptotic signaling. PLoS Pathog 19, e1011297 (2023).

65. E. A. Wall, et al., Suppression of LPS-Induced TNF-α Production in Macrophages by cAMP Is Mediated by PKA-AKAP95-p105. Science Signaling 2, ra28–ra28 (2009).

66. K. Rohde, R. M. Yates, G. E. Purdy, D. G. Russell, Mycobacterium tuberculosis and the environment within the phagosome. Immunological Reviews 219, 37–54 (2007).

67. A. K. Pandey, C. M. Sassetti, Mycobacterial persistence requires the utilization of host cholesterol. Proc Natl Acad Sci U S A 105, 4376–4380 (2008).

68. N. V. Simwela, E. Jaecklein, C. M. Sassetti, D. G. Russell, Impaired fatty acid import or catabolism in macrophages restricts intracellular growth of Mycobacterium tuberculosis. eLife 13, RP102980 (2025).

69. C. Toniolo, N. Dhar, J. D. McKinney, Uptake-independent killing of macrophages by extracellular Mycobacterium tuberculosis aggregates. The EMBO Journal 42, e113490 (2023).

70. C. K. Stover, et al., New use of BCG for recombinant vaccines. Nature 351, 456–460 (1991).

71. C. Toniolo, D. Sage, J. D. McKinney, N. Dhar, “Quantification of Mycobacterium tuberculosis Growth in Cell-Based Infection Assays by Time-Lapse Fluorescence Microscopy” in Intracellular Pathogens: Methods and Protocols, A. Thakur, Ed. (Springer US, 2024), pp. 167–188.

72. B. Bosch, et al., Genome-wide gene expression tuning reveals diverse vulnerabilities of M. tuberculosis. Cell 184, 4579–4592.e24 (2021).

73. J. Schindelin, et al., Fiji: an open-source platform for biological-image analysis. Nat Methods 9, 676–682 (2012).

74. D. R. Stirling, et al., CellProfiler 4: improvements in speed, utility and usability. BMC Bioinformatics 22, 433 (2021).

75. A. Ducret, E. M. Quardokus, Y. V. Brun, MicrobeJ, a tool for high throughput bacterial cell detection and quantitative analysis. Nat Microbiol 1, 16077 (2016).

76. C. Vilchèze, et al., Rational Design of Biosafety Level 2-Approved, Multidrug-Resistant Strains of Mycobacterium tuberculosis through Nutrient Auxotrophy. mBio 9, 10.1128/mbio.00938-18 (2018).

