## Supplementary Data for "A genetically encoded biosensor reveals heterogenous cAMP dynamics coordinating growth and resuscitation in Mycobacterium tuberculosis"

Shaun Wachter *et al.*

**This PDF file includes:**

Figs. S1 to S12
Tables S1 to S2
Movie Legends S1 to S4

**Other Supplementary Materials for this manuscript include the following:**

Movies S1 to S4

Supplementary Figure 1

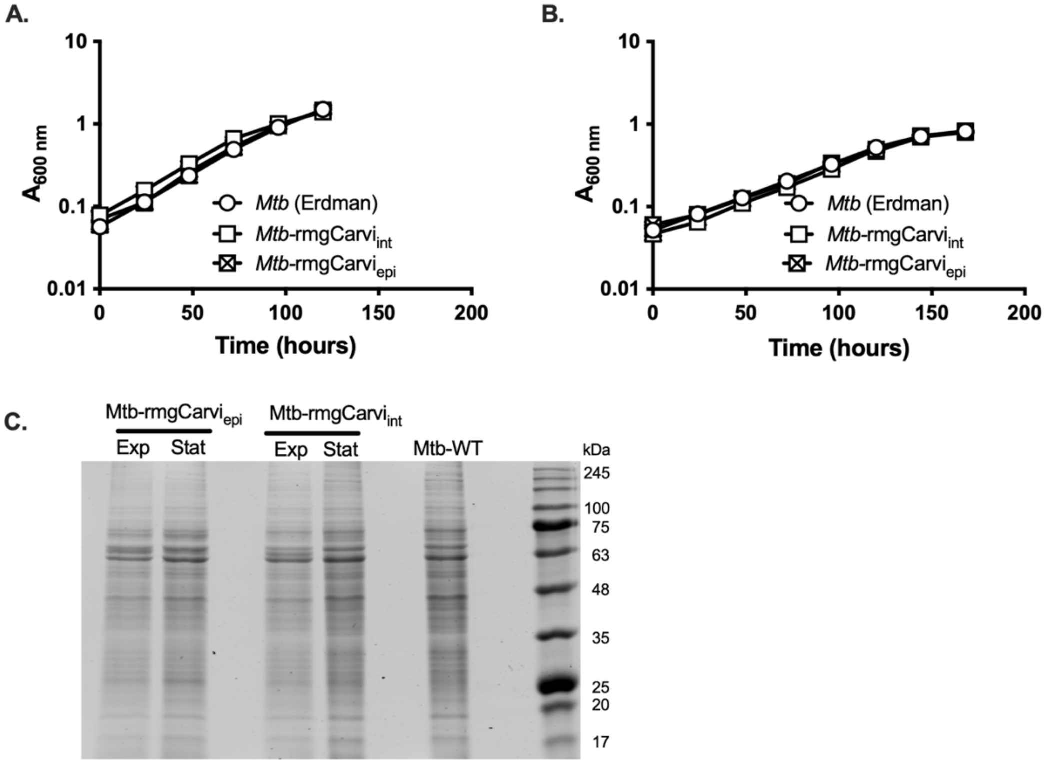

**Supplementary Figure 1. Expression of rmgCarvi does not alter growth kinetics of *M. tuberculosis*.** (**A, B**) Growth curves of Mtb-rmgCarvi$_{int}$ (open squares), Mtb-rmgCarvi$_{epi}$ (crossed-out squares), compared to wild-type Mtb (Erdman) (circles), in 7H9+OADC media (**A**), and in growth media containing butyric acid, palmitic acid, and cholesterol (BCP) as the sole carbon sources (**B**). Data represents the mean ± SD of three biological replicates. (**C**) Crude cell-free extracts of Mtb transformed with rmgCarvi$_{epi}$ or rmgCarvi$_{int}$ were analysed on a 0.1% SDS-10% PAGE and stained with Coomassie Brilliant Blue. The molecular weight markers are indicated on the right. Samples were collected from cultures in exponential (Exp) and stationary phase (Stat).

**Supplementary Figure 2**

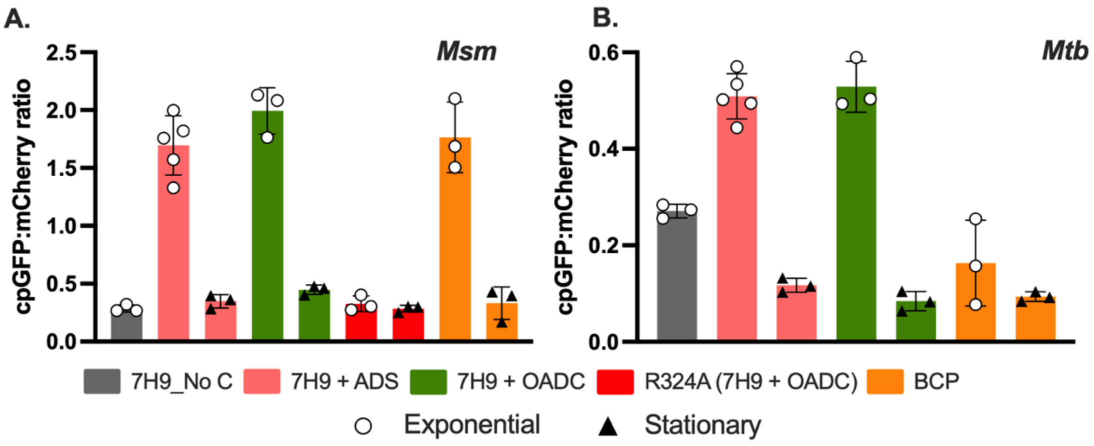

**Supplementary Figure 2. Cyclic-AMP levels decline during stationary phase in *M. smegmatis* and *M. tuberculosis*. (A, B)** Msm-rmgCarvi$_{epi}$ or Mtb-rmgCarvi$_{epi}$ were cultured in various growth media – basal 7H9 media with no added carbon source (7H9_No C, gray); 7H9+ADS (pink); 7H9+OADC (green) or cholesterol and fatty-acid-based BCP media (orange). Fluorescence (cpGFP and mCherry) was measured from cultures in exponential (open circles) or stationary (solid triangles) phase, using a plate reader. Ratiometric rmgCarvi fluorescence values reveal a substantial reduction in cAMP levels during stationary phase relative to exponential growth in both Msm and Mtb. Introduction of the R324A mutation in rmgCarvi's cAMP-binding domain (CBD) abolished the biosensor's ability to respond to cAMP. Data represents the mean ± SD from 3-5 independent experiments.

Supplementary Figure 3

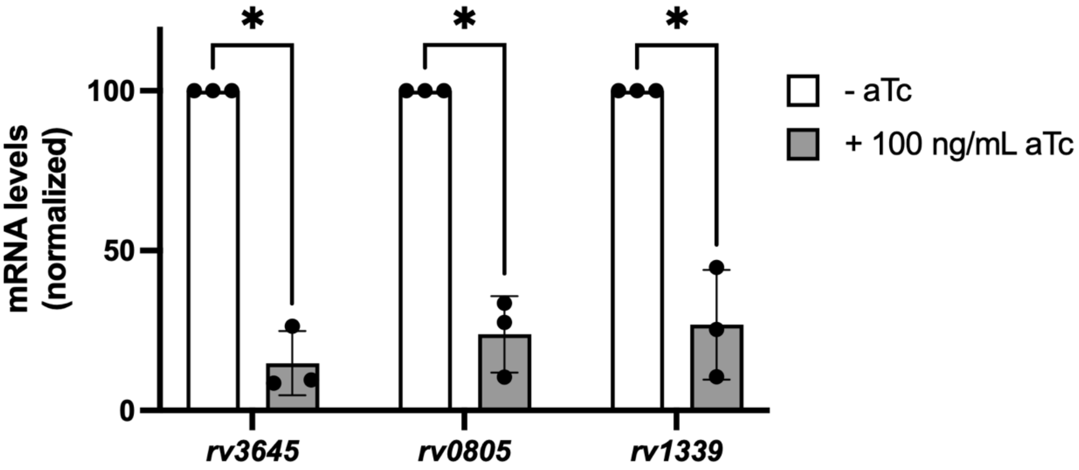

**Supplementary Figure 3. CRISPRi mediated repression of cAMP-related genes.** Early exponential cultures of Mtb-rmgCarvi$_{epi}$ (mc$^{2}$6206) strains expressing sgRNAs targeting *rv3645* or *rv0805* or *rv1339* were induced with 100 ng/mL anhydrotetracycline (aTc; grey bars) for 48-hours or left uninduced (white bars). Expression of targeted genes was determined by qRT-PCR, indicating a ~75-85% reduction in expression of the targeted genes upon CRISPRi-mediated gene knockdown. Results are the mean ± SD of three biological replicates. Gene expression in the uninduced samples was normalized to 100% and percent expression upon aTc induction was calculated. Unpaired t-test, * - $p < 0.02$.

Supplementary Figure 4

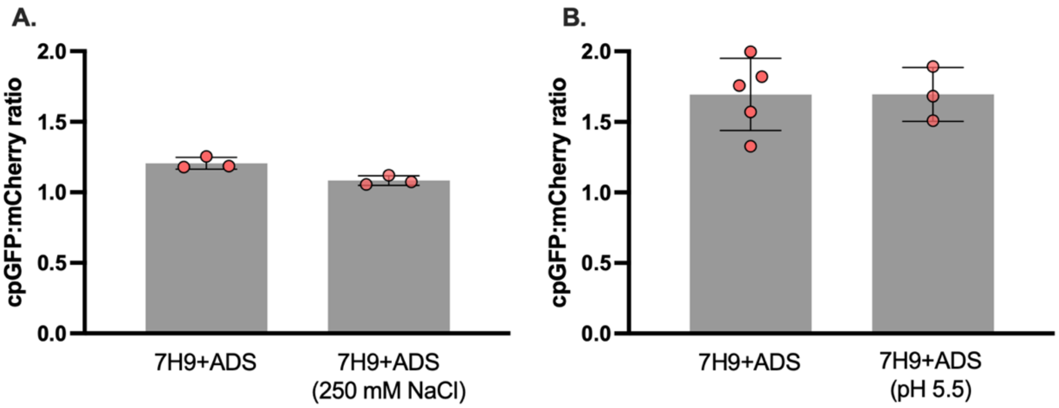

**Supplementary Figure 4. Cyclic-AMP levels in *M. smegmatis* are unaffected by salt and acid stress. (A, B)** Msm-rmgCarvi$_{epi}$ was cultured in 7H9+ADS and exposed to high concentrations of salt (250 mM NaCl ; **A**) or acidic stress (pH 5.5; **B**), for a duration of 1h. Ratiometric rmgCarvi fluorescence, measured using a plate reader, showed no change in intramycobacterial cAMP levels under either condition. Data represents the mean ± SD from 3-5 biological replicates.

**Supplementary Figure 5**

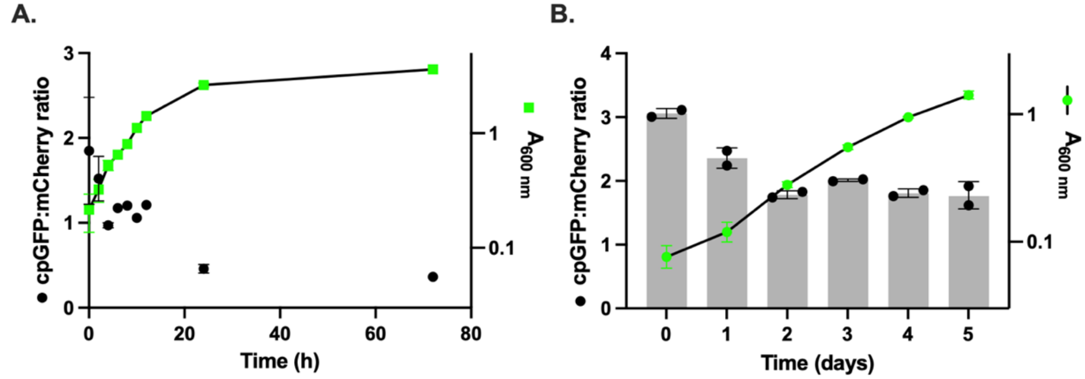

**Supplementary Figure 5. Mycobacterial cyclic-AMP levels decline as cultures age.** (**A**) Plot depicts the ratiometric rmgCarvi fluorescence (left *y-axis, black circles*) and $A_{600\ nm}$ (right *y-axis, green squares*) of Msm-rmgCarvi$_{epi}$, measured using a plate reader. N = 2-3. **(B)** Plot depicts the ratiometric rmgCarvi fluorescence (left *y-axis, gray bars and black circles*) and $A_{600\ nm}$ (right *y-axis, green circles*) of Mtb-rmgCarvi$_{epi}$ ($H_{37}Rv$), at the indicated time points, measured by flow cytometry. N = 2. Both strains were cultured in 7H9+ADS. Results shown are mean ± SD.

Supplementary Figure 6

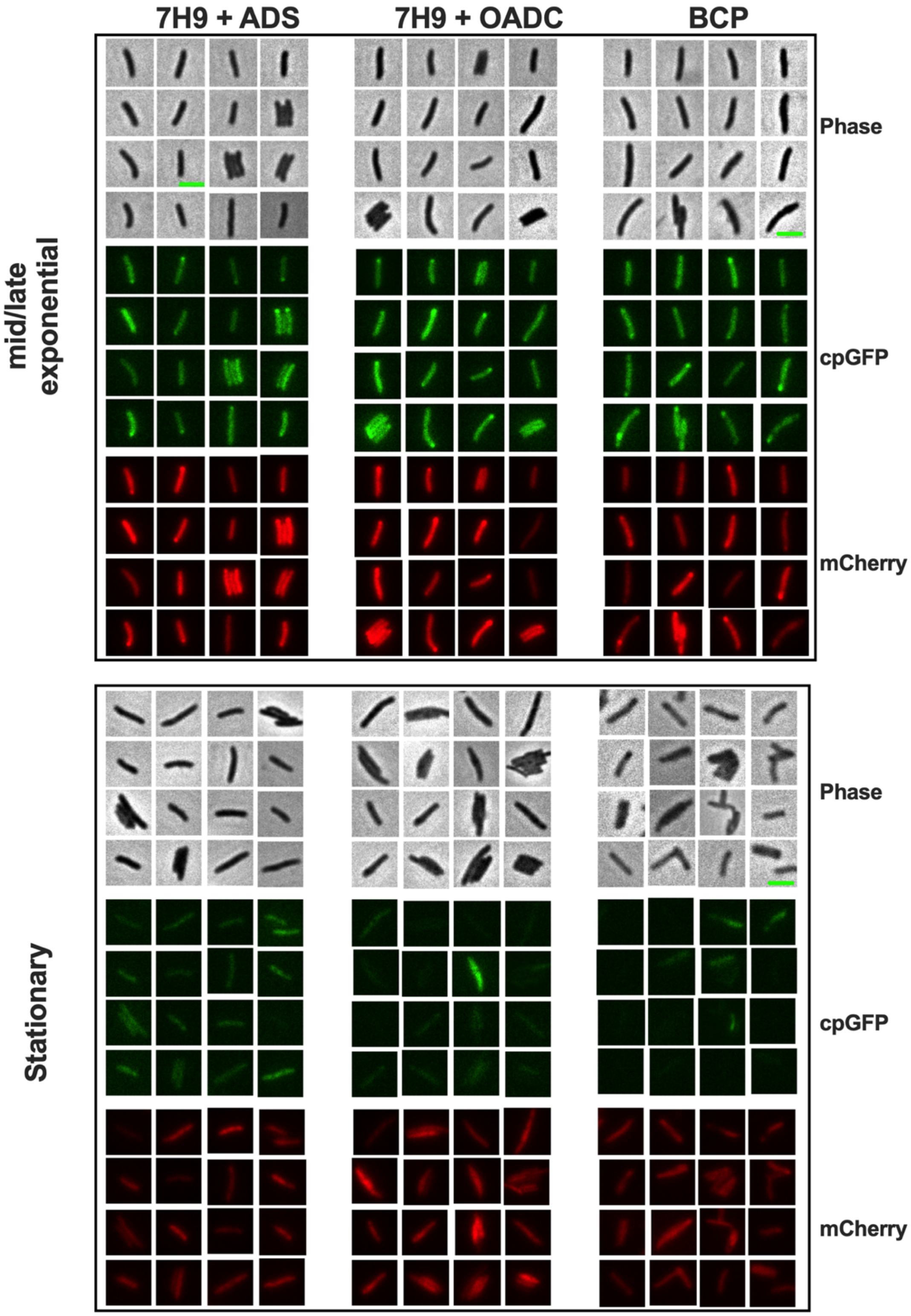

**Supplementary Figure 6. Single-cell imaging of *M. tuberculosis* expressing rmgCarvi.** Representative snapshots of Mtb-rmgCarvi$_{epi}$ (Erdman) bacteria cultured in 7H9+ADS or 7H9+OADC or BCP media, imaged during the different phases of growth. Bacteria were spotted on agarose pads and images acquired on phase, cpGFP (Ex$_{475\ nm}$/Em$_{535\ nm}$) and mCherry (Ex$_{575\ nm}$/Em$_{642\ nm}$) channels using a 100X objective. The green scale bar represents 3 µm.

**Supplementary Figure 7**

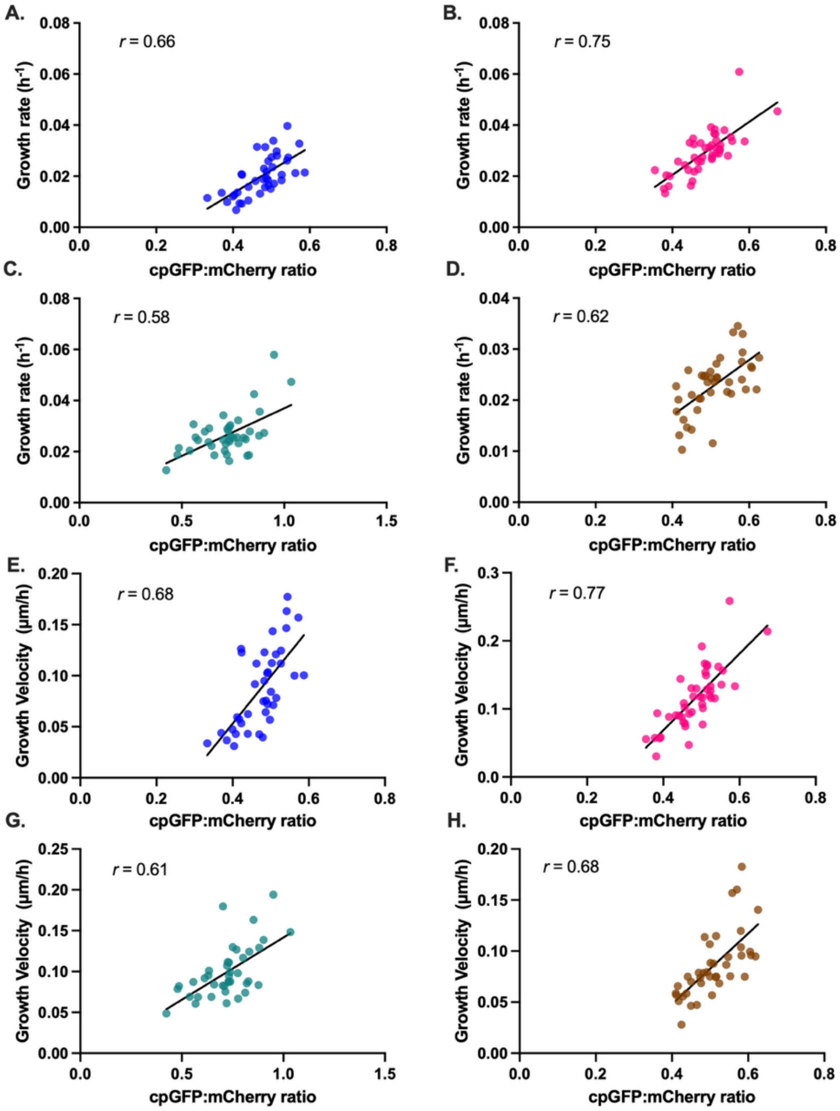

**Supplementary Figure 7. Cyclic-AMP levels in *M. tuberculosis* are reflective of culture condition and strongly correlated to growth rate and velocity. (A – H)** Mtb-rmgCarvi$_{int}$ (Erdman) cultured in microfluidic device in 7H9+ADS were imaged every hour on phase, cpGFP (Ex$_{475\ nm}$/Em$_{535\ nm}$) and mCherry (Ex$_{575\ nm}$/Em$_{642\ nm}$) channels using a 100X objective lens. Each of the four plots depicts the correlation of average ratiometric rmgCarvi fluorescence (*x-axis*) with growth rate (*y-axis*) (**A-D**) or growth velocity (*y-axis*) (**E-H**) in four independent biological replicates (N=40-42 in each replicate). Pearson correlation coefficient *(r)* values are shown, $p <$ 0.0001, for each of the correlations.

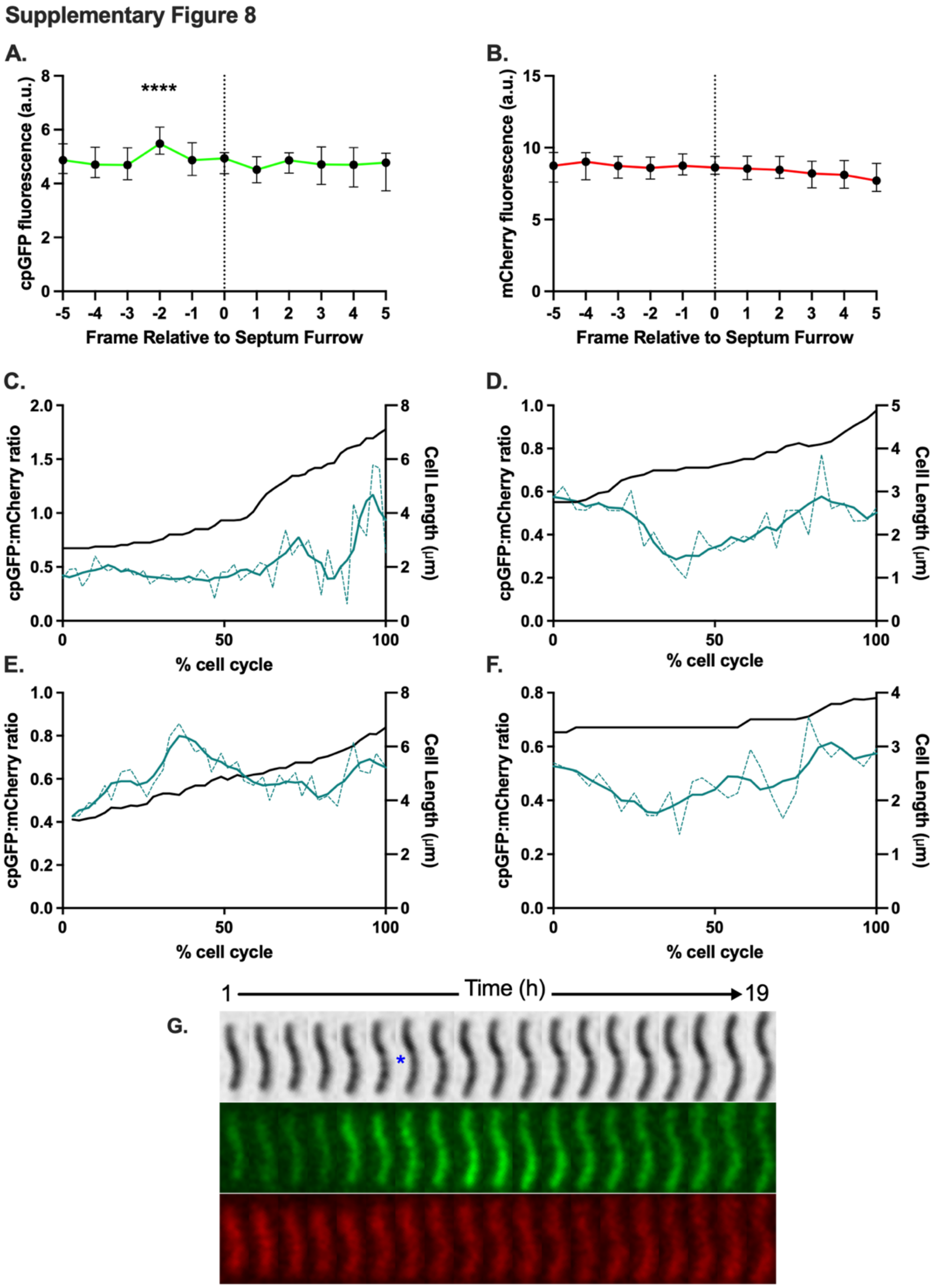

**Supplementary Figure 8. Spike in rmgCarvi fluorescence prior to cell division in *M. tuberculosis* is restricted to the cpGFP channel.** (**A, B**) Median cpGFP (**A**) and mCherry (**B**) fluorescence in Mtb-rmgCarvi$_{int}$ (Erdman) grown in a microfluidic device, plotted for five frames before and five frames after septum furrow formation seen on the phase channel. N=81, pooled from three independent experiments. The pre-division fluorescence spike observed in the rmgCarvi fluorescence ratio (**Fig. 4D, E**) is driven exclusively by a transient increase in cpGFP signal (**A**) and is not reflected in the mCherry signal (**B**), indicating a genuine increase in cAMP rather than a fluorescence artifact. Data represents medians ± 95% CI. **** - $p < 0.0001$, One-way ANOVA with Geisser-Greenhouse and Holm-Sidak's corrections. (**C-F**) Plots depict the ratiometric rmgCarvi fluorescence (solid green lines – smoothed data, dashed green lines – raw data, left *y-axis*) and cell-length (black lines, right *y-axis*) of representative Mtb-rmgCarvi$_{int}$ bacteria, mapped from their birth to division (*x-axis*). (**G**) Representative image sequence of an Mtb-rmgCarvi$_{int}$ bacterial cell before, and after cell division, depicting the cAMP pulsing event. The appearance of the furrow in the septal region is indicated with a blue asterisk. Bacteria were cultured in a microfluidic device and images acquired on phase (top), cpGFP (Ex$_{475\ nm}$/Em$_{535\ nm}$, middle) and mCherry (Ex$_{575\ nm}$/Em$_{642\ nm}$, bottom) channels using a 100X objective. The numbers above indicate the time span of the image series.

### Supplementary Figure 9

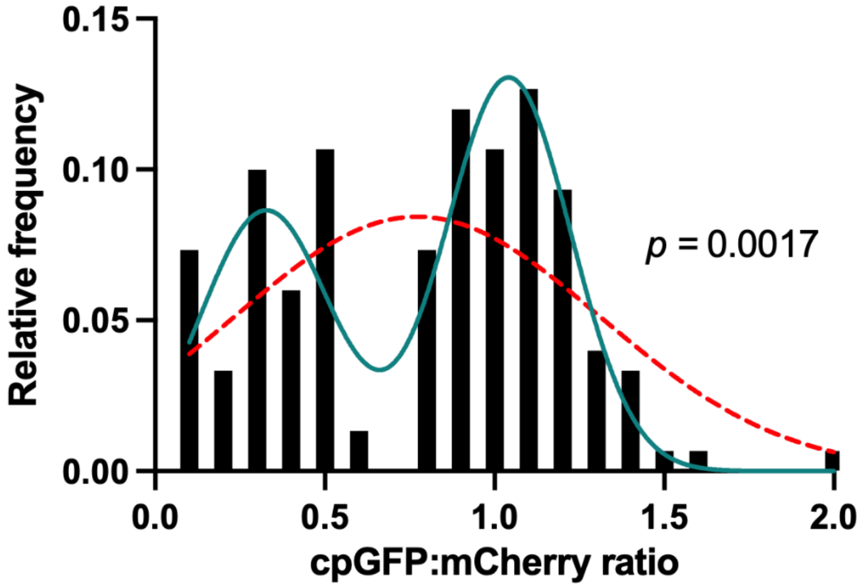

| | Goodness of fit | | Probability model is correct |
|---|---|---|---|
| | $R^2$ | AICc | |
| **Gaussian** | 0.3959 | -123.5 | 1.52% |
| **Sum of two Gaussians** | 0.7887 | -131.8 | 98.48% |

**Supplementary Figure 9. Distribution of ratiometric rmgCarvi fluorescence in nutrient-starved *M. tuberculosis* is bimodal.** Histogram plot of rmgCarvi fluorescence ratios averaged over five hours from Mtb-rmgCarvi$_{int}$ (Erdman) cultures starved in PBS+0.02% tyloxapol for 5-6 weeks, overlaid with GraphPad Prism fits: unimodal "Gaussian (red dotted line) and bimodal "Sum of two Gaussians" (teal solid line). The table below lists the fit statistics comparing the two models - goodness of fit ($R^2$), corrected Akaike information criterion (AICc), and relative probability of model correctness. Values indicate that the sum of two Gaussians bimodal model provides a substantially better fit than a single Gaussian, consistent with phenotypic bifurcation of Mtb during prolonged nutrient starvation. N=150.

Supplementary Figure 10

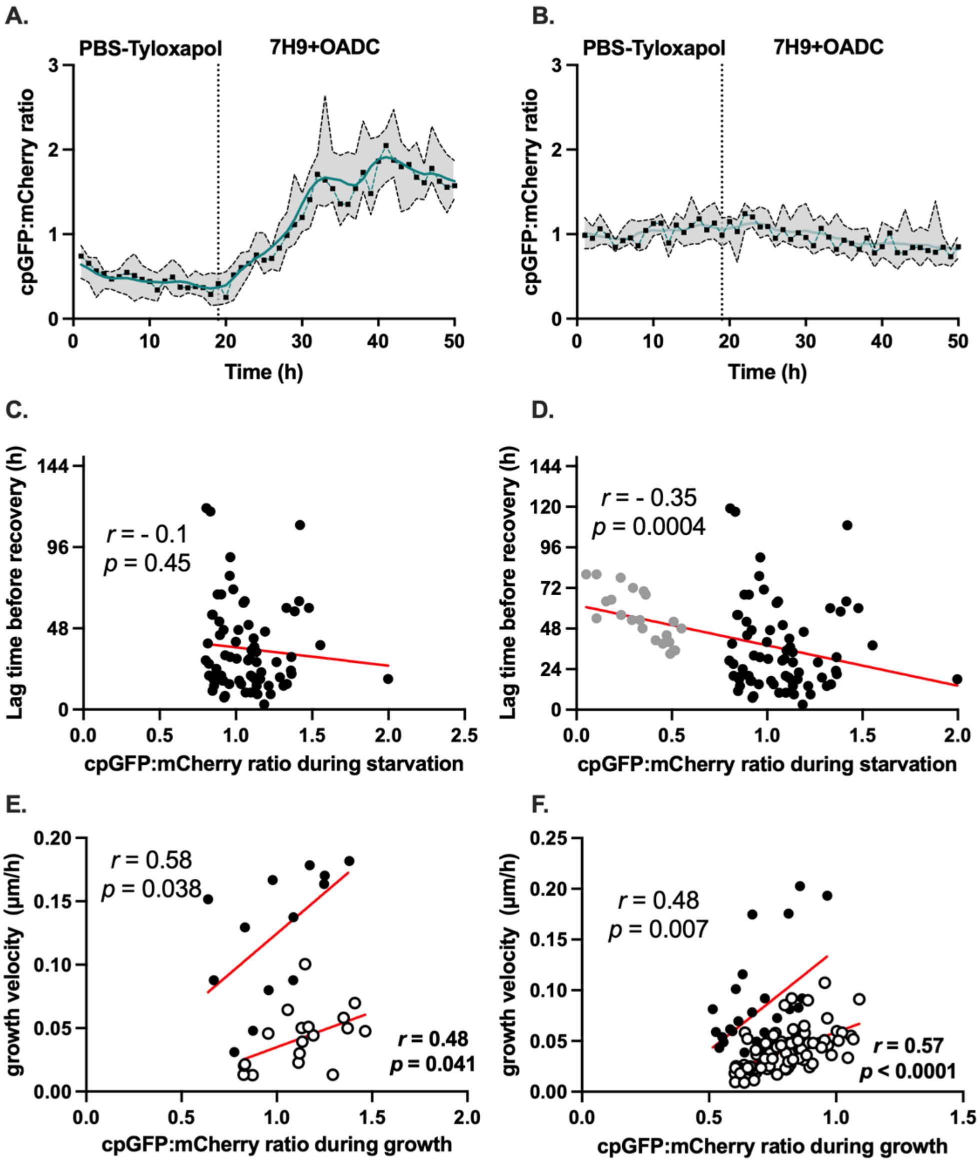

**Supplementary Figure 10. Cyclic-AMP-driven hysteresis influences regrowth dynamics in starved *M. tuberculosis*.** (**A, B**) Mtb-rmgCarvi$_{int}$ (Erdman) cultures starved in PBS+0.02% tyloxapol for 5-6 weeks were seeded into microfluidic device and exposed to fresh media (7H9+OADC). Line traces represent the median ratiometric rmgCarvi fluorescence (black squares - raw data; green line – smoothed data) in cells originating from low-cAMP (**A**) and high-cAMP (**B**) subpopulations (N=8, for each), measured by time-lapse microscopy. The gray shaded area represents the interquartiles of the median. (**C, D**) Correlation plots between ratiometric rmgCarvi fluorescence and the lag time before initiation of growth upon nutrient supplementation, in high cAMP subpopulation (**C**, N=71) and the combined low and high-cAMP subpopulations of Mtb (**D**, N=93). Pearson correlation coefficients (*r*) and the associated *p* values are shown. The red line represents linear fit to the data. (**E,F**) Correlation plots of ratiometric rmgCarvi fluorescence (cAMP content) during regrowth from starvation and growth velocity in "low" cAMP cells from two independent biological replicates, black circles (N=13 cells, Pearson correlation coefficient, $r = 0.58$, $p = 0.038$) and white circles (N=18 cells, Pearson correlation coefficient, $r = 0.48$, $p = 0.041$) (**E**) and in "high" cAMP cells from two independent biological replicates, black circles (N=30 cells, Pearson correlation coefficient, $r = 0.48$, $p = 0.007$) and white circles (N=83 cells, Pearson correlation coefficient, $r = 0.57$, $p < 0.0001$) (**F**).

**Supplementary Figure 11**

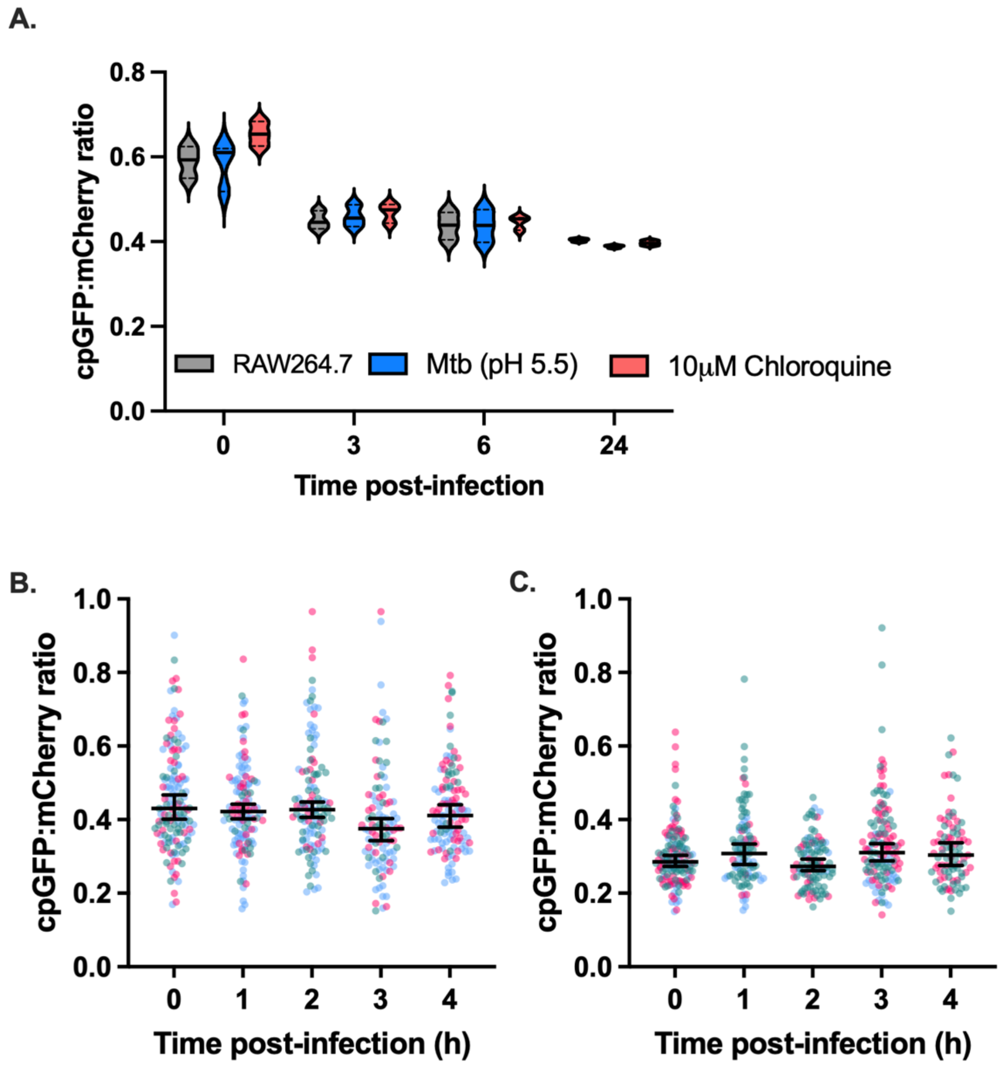

**Supplementary Figure 11.** (**A**) **Phagosomal pH alone does not induce mycobacterial cAMP during infection.** rmgCarvi fluorescence dynamics in Mtb-rmgCarvi$_{epi}$ during the first 24 hours of infection of RAW264.7 macrophages, or Mtb-rmgCarvi$_{epi}$ pre-cultured in acidic pH (pH 5.5) for 3 hours prior to infection and in macrophages whose phagosomal pH was deacidified using 10 µM chloroquine (treated for 24 hours prior to infection), measured by flow cytometry. Intra-mycobacterial cAMP levels drop within 3 hours and is maintained stably over the next 21 hours. cpGFP and mCherry fluorescence was measured using flow cytometry (N=3, 10000 cells per replicate per time point). (**B, C**) **rmgCarvi fluorescence in extracellular *M. tuberculosis* remains unchanged during macrophage infection.** Quantification of rmgCarvi fluorescence ratios in extracellular Mtb-rmgCarvi$_{epi}$ bacteria during infection of murine RAW264.7 (**B**) or human U937 (**C**) macrophages. Images were analyzed in FIJI; N=3 biological replicates with >100 extracellular bacteria per replicate per time point. No significant change in ratiometric rmgCarvi fluorescence was observed over the infection time course. Data represents medians ± 95% CI.

**Supplementary Figure 12**

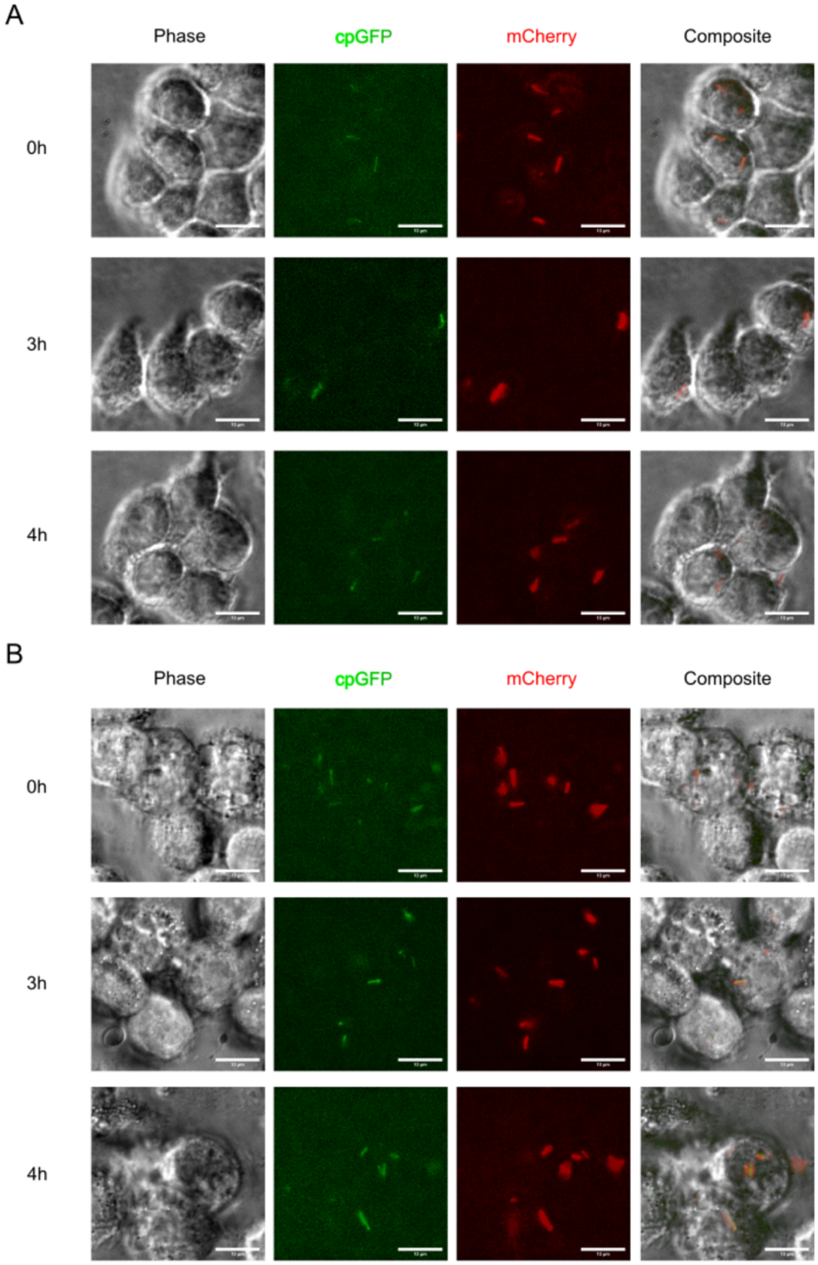

**Supplementary Figure 12. Representative images of macrophages infected with *M. tuberculosis* expressing rmgCarvi.** (**A, B**) Representative fluorescence microscopy images of RAW264.7 murine macrophages (**A**) and PMA-differentiated U937 human macrophages (**B**) infected with Mtb-rmgCarvi$_{epi}$ (Erdman). Images were acquired on phase, cpGFP (Ex$_{475\ nm}$/Em$_{535\ nm}$) and mCherry (Ex$_{575\ nm}$/Em$_{642\ nm}$) channels using a 100X objective. Composite channel represents the merged composites at 0-, 3-, and 4-hours post-infection. Scale bar = 10 µm.

**Table S1. Bacterial strains, primers and plasmids used in this study**

| **Name** | **Description** | **Reference** |
|---|---|---|
| **Strains** | | |
| *Mycobacterium smegmatis* (Msm) | *Mycobacterium smegmatis* mc$^2$155, wild-type strain | Lab collection |
| *Mycobacterium tuberculosis* (auxotroph) | *Mycobacterium tuberculosis* $H_{37}$Rv (Δ*panCD* Δ*leuCD*) mc$^2$6206, wild-type strain | Bill Jacobs Lab. Vilchèze *et al*, 2018 (76) |
| *Mycobacterium tuberculosis* (Erdman) | *Mycobacterium tuberculosis* Erdman, wild-type strain | Lab collection |
| *Mycobacterium tuberculosis* ($H_{37}$Rv) | *Mycobacterium tuberculosis* $H_{37}$Rv, wild-type strain | Lab collection |
| Msm-rmgCarvi$_{int}$/ Mtb-rmgCarvi$_{int}$ | Mycobacterial parental strain transformed with pND257-rmgCarvi plasmid | This study |
| Msm-rmgCarvi$_{epi}$/ Mtb-rmgCarvi$_{epi}$ | Mycobacterial parental strain transformed with pMV261-rmgCarvi plasmid | This study |
| Mtb-rmgCarvi$_{epi}$-CRISPRi-Rv1339 | Mtb-rmgCarvi$_{epi}$ (mc$^2$6206) transformed with pIRL2-Hyg-sgRNA-Rv1339 | This study |
| Mtb-rmgCarvi$_{epi}$-CRISPRi-Rv0805 | Mtb-rmgCarvi$_{epi}$ (mc$^2$6206) transformed with pIRL2-Hyg-sgRNA-Rv0805 | This study |
| Mtb-rmgCarvi$_{epi}$-CRISPRi-Rv3645 | Mtb-rmgCarvi$_{epi}$ (mc$^2$6206) transformed with pIRL2-Hyg-sgRNA-Rv3645 | This study |
| Msm-rmgCarvi$_{epi}$-R324A | *Mycobacterium smegmatis* mc$^2$155 transformed with pMV261-rmgCarvi-R324A plasmid | This study |
| **Primers** | | |
| **Name** | **Sequence (5' - 3')** | **Reference** |
| rmgCarvi-hsp60-NheI-F | ACG TGC TAG CAT GGC CAA GAC AAT TGC GGT GAG CAA GGG CGA GG | This study |
| gCarvi-ScaI-R | ACG TAG TAC TTC AGT TGT ACT CCA GCT TGT G | This study |
| pND257_Flank_Seq_F | CCG AAA TGA GCA CGA TCC GC | This study |
| pND257_Flank_Seq_R | GCC AGG AGC ATT GCC GTT C | This study |
| rmgCarvi_Internal_Seq_F | CAA CGA GGA CTA CAC CAT CGT G | This study |
| pIRL2-Seq-F | TTC CTG TGA AGA GCC ATT GAT AAT G | This study |
| rmgCarvi_Q5_R324A_F | AGG ACA GGA GGC GTC CGC TTG GG | This study |
| rmgCarvi_Q5_R324A_R | TCC TCG AAC AGG CCC | This study |

| | | |
|---|---|---|
| Rv1339_sgRNA_1_F | GGG AAC TGT GCC ATG CCC GAT ACA ACG AT | This study |
| Rv1339_sgRNA_1_R | AAA CAT CGT TGT ATC GGG CAT GGC ACA GT | This study |
| Rv3645_sgRNA_1_F | GGG AAC CGT ATC GGG CGG CGG GGA CTT G | This study |
| Rv3645-sgRNA_1_R | AAA CCA AGT CCC CGC CGC CCG ATA CGG T | This study |
| Rv0805_sgRNA_1_F | GGG AGA GAT GAG TGT CGC TGA TAT GTA | This study |
| Rv0805_sgRNA_1_R | AAA CTA CAT ATC AGC GAC ACT CAT CTC | This study |
| Rv1339-qRT-F | CAT GGC ACA GTA GTG TCC GTG | This study |
| Rv1339-qRT-R | CAC ATG CAC CGA CGC G | This study |
| Rv3645-qRT-F | ACA TCA TCG GCG CAT TGT | This study |
| Rv3645-qRT-R | GCT GAC GAA GAT CAG TAG GTT G | This study |
| Rv0805-qRT-F | ATC AGC GAC ACT CAT CTC ATC | This study |
| Rv0805-qRT-R | AAG GCC GGA TTG GTT CAA | This study |
| Mtb_SigA_qRT_F | AAA CAG ATC GGC AAG GTA GC | This study |
| Mtb_SigA_qRT_R | CTG GAT CAG GTC GAG AAA CG | This study |
| Hyg-pIRL2-NheI-F | ACG TGC TAG CGT TTT AAA TCA ATC TAA AGT ATA TAT GAG TAA ACT TGG TC | This study |
| Hyg-pIRL2-EcoRV-R | ACG TGA TAT CTC AGG CGC CGG G | This study |
| **Plasmids** | | |
| **Name** | **Purpose** | **Reference** |
| pCR2.1-TOPO | Cloning plasmid, Amp$^r$, Kan$^r$ | Invitrogen |
| pND257 | Mycobacterial L5-based integrative plasmid. Kan$^r$ | Toniolo *et al*, 2023 (69) |
| pMV261 | Mycobacterial episomal plasmid. Kan$^r$ | Stover *et al*, 1991 (70) |
| pND257-rmgCarvi | Mycobacterial L5-based integrative plasmid encoding rmgCarvi under the control of strong UV15 promoter, Kan$^r$ | This study |
| pMV261-rmgCarvi | Mycobacterial episomal plasmid encoding rmgCarvi under the control of strong strong UV15 promoter, Kan$^r$ | This study |
| pMV261-rmgCarvi-R324A | Mycobacterial episomal plasmid encoding cAMP binding-deficient rmgCarvi under the control of strong strong UV15 promoter, Kan$^r$ | This study |
| pIRL2 | Mycobacterial L5-based integrative plasmid. Encodes all CRISPRi machinery necessary for functionality in *Mycobacterium spp*. No sgRNA sequence, Kan$^r$ | Bosch *et al*, 2021 (72) |

| | | |
|---|---|---|
| pIRL2-Hyg | Mycobacterial L5-based integrative plasmid. Encodes all CRISPRi machinery necessary for functionality in Mycobacterium spp. No sgRNA sequence, $Hyg^r$ | This study |
| pIRL2-Hyg-sgRNA-Rv1339 | pIRL2-Hyg plasmid encoding Rv1339 sgRNA sequence, $Hyg^r$ | This study |
| pIRL2-Hyg-sgRNA-Rv0805 | pIRL2-Hyg plasmid encoding Rv0805 sgRNA sequence, $Hyg^r$ | This study |
| pIRL2-Hyg-sgRNA-Rv3645 | pIRL2-Hyg plasmid encoding Rv3645 sgRNA sequence. $Hyg^r$ | This study |

$Amp^r$ – Ampicillin resistant; $Hyg^r$ – Hygromycin resistant; $Kan^r$ – Kanamycin resistant.

**Table S2: Descriptive statistics of *M. tuberculosis*-rmgCarvi cells cultured in different growth medium and imaged at different phases of growth.**

**Growth medium (7H9+ADS)**

| | Early-exponential ($A_{600\ nm}$ 0.1-0.3) | Mid-exponential ($A_{600\ nm}$ 0.3-0.5) | Late-exponential ($A_{600\ nm}$ 0.5-0.8) | Stationary phase ($A_{600\ nm}$ >1.0) |
|---|---|---|---|---|
| **Mean** | 0.1592 | 0.1134 | 0.2135 | 0.0944 |
| **SD** | 0.0560 | 0.0511 | 0.0493 | 0.076 |
| **Median** | 0.1559 | 0.1098 | 0.2064 | 0.0701 |
| **95% CI of median (Lower, Upper)** | (0.15, 0.1591) | (0.1065, 0.1152) | (0.2017, 0.2124) | (0.0619, 0.0814) |
| **$CV^2$** | 0.1238 | 0.2033 | 0.0534 | 0.6480 |
| **N** | 507 | 373 | 470 | 446 |

**Growth medium (7H9+OADC)**

| | Early-exponential ($A_{600\ nm}$ 0.1-0.3) | Mid-exponential ($A_{600\ nm}$ 0.3-0.5) | Late-exponential ($A_{600\ nm}$ 0.5-0.8) | Stationary phase ($A_{600\ nm}$ >1.0) |
|---|---|---|---|---|
| **Mean** | 0.2209 | 0.0955 | 0.2064 | 0.0627 |
| **SD** | 0.0587 | 0.0545 | 0.0568 | 0.0607 |
| **Median** | 0.2244 | 0.0888 | 0.1961 | 0.0412 |
| **95% CI of median (Lower, Upper)** | (0.2198, 0.2327) | (0.0835, 0.0941) | (0.189, 0.2016) | (0.0385, 0.045) |
| **$CV^2$** | 0.0707 | 0.3265 | 0.076 | 0.9388 |
| **N** | 434 | 415 | 443 | 629 |

**Growth medium (BCP)**

| | Early-exponential ($A_{600\ nm}$ 0.1-0.3) | Mid-exponential ($A_{600\ nm}$ 0.3-0.5) | Late-exponential ($A_{600\ nm}$ 0.5-0.8) | Stationary phase ($A_{600\ nm}$ >1.0) |
|---|---|---|---|---|
| **Mean** | 0.0833 | 0.199 | 0.2021 | 0.062 |
| **SD** | 0.0468 | 0.037 | 0.0518 | 0.0552 |
| **Median** | 0.0737 | 0.1960 | 0.1939 | 0.0439 |
| **95% CI of median (Lower, Upper)** | (0.0661, 0.078) | (0.1893, 0.1999) | (0.1873, 0.1999) | (0.0398, 0.0498) |
| **$CV^2$** | 0.3163 | 0.0346 | 0.0658 | 0.7926 |
| **N** | 336 | 332 | 442 | 200 |

### Supplementary Movie Legends

**Supplementary Movie S1. Timelapse microscopy of *M. tuberculosis* (Erdman) expressing rmgCarvi$_{int}$ (first example).** Representative timelapse microscopy experiment in which Mtb-rmgCarvi$_{int}$ was cultured in a microfluidic device and imaged for ~ 60 hours under a constant flow of 7H9+ADS. Images were acquired every hour on phase (left, gray), cpGFP (middle, green) and mCherry (right, magenta) channels. The spike in fluorescence is clearly visible on the cpGFP channel around the time the cell divides. Numbers (upper right corner) indicate hours elapsed and the scale bar on bottom left, represents 3 μm.

**Supplementary Movie S2. Timelapse microscopy of *M. tuberculosis* (Erdman) expressing rmgCarvi$_{int}$ (second example).** Representative timelapse microscopy experiment in which Mtb-rmgCarvi$_{int}$ was cultured in a microfluidic device and imaged for ~ 60 hours under a constant flow of 7H9+ADS. Images were acquired every hour on phase (left, gray), cpGFP (middle, green) and mCherry (right, magenta) channels. The spike in fluorescence is clearly visible on the cpGFP channel around the time the cell divides. Numbers (upper right corner) indicate hours elapsed and the scale bar on bottom left, represents 3 μm.

**Supplementary Movie S3. Timelapse microscopy showing resuscitation of starved *M. tuberculosis* (Erdman) expressing rmgCarvi$_{int}$ (low cAMP subpopulation).** Three representative timelapse microscopy image series in which Mtb-rmgCarvi$_{int}$ bacteria (starved for 6-8 weeks) were seeded in a microfluidic device and imaged for ~ 18 hours in starvation medium (PBS+0.02% tyloxapol) before being exposed to 7H9+OADC medium for the next 150 hours. Images were acquired every hour on phase (left, gray), cpGFP (middle, green) and mCherry (right, magenta) channels, and only the time period 0-115 h is shown. This subset of cells have low cpGFP fluorescence during starvation, and after a significant lag following nutrient exposure, induce cpGFP and commence elongation. Numbers (upper right corner) indicate hours elapsed and the scale bar on bottom left, represents 3 μm. The media condition is indicated in the upper middle region (PBS/7H9).

**Supplementary Movie S4. Timelapse microscopy showing resuscitation of starved *M. tuberculosis* (Erdman) expressing rmgCarvi$_{int}$ (high cAMP subpopulation).** Three representative timelapse microscopy image series in which Mtb-rmgCarvi$_{int}$ bacteria (starved for 6-8 weeks) were seeded in a microfluidic device and imaged for ~ 18 hours in starvation medium (PBS+0.02% tyloxapol) before being exposed to 7H9+OADC medium for the next 150 hours.

Images were acquired every hour on phase (left, gray), cpGFP (middle, green) and mCherry (right, magenta) channels, and only the time period 0-115 h is shown. This cell subset exhibits high cpGFP fluorescence during starvation, and upon nutrient exposure, begins elongation accompanied by a decrease in cpGFP fluorescence. Numbers (upper right corner) indicate hours elapsed and the scale bar on bottom left, represents 3 μm. The media condition is indicated in the upper middle region (PBS/7H9).
